# Multiparametric senescent cell phenotyping reveals CD24 osteolineage cells as targets of senolytic therapy in the aged murine skeleton

**DOI:** 10.1101/2023.01.12.523760

**Authors:** Madison L. Doolittle, Dominik Saul, Japneet Kaur, Jennifer L. Rowsey, Stephanie J. Vos, Kevin D. Pavelko, Joshua N. Farr, David G. Monroe, Sundeep Khosla

**Affiliations:** Division of Endocrinology, Diabetes and Metabolism, Mayo Clinic, Rochester, MN, 55905, USA; Robert and Arlene Kogod Center on Aging, Mayo Clinic, Rochester, MN 55905, USA; Department for Trauma and Reconstructive Surgery, BG Clinic, University of Tübingen, Germany; Department of Immunology, Mayo Clinic, Rochester, MN, 55905, USA

**Author notes:** **Correspondence:** Sundeep Khosla, M.D., Guggenheim 7-11, Mayo Clinic College of Medicine, 200 First Street SW, Rochester, MN 55905.

**Keywords:** skeleton, osteolineage, osteoblasts, osteocytes, aging, senescence, SASP, mass cytometry

## Abstract

Senescence drives organismal aging, yet the deep characterization of senescent cells *in vivo* remains incomplete. Here, we applied mass cytometry by time-of-flight (CyTOF) using carefully validated antibodies to analyze senescent cells at single-cell resolution. We used multiple criteria to identify senescent mesenchymal cells that were growth arrested and resistant to apoptosis (p16+/Ki67-/BCL-2+; “p16KB” cells). These cells were highly enriched for senescence-associated secretory phenotype (SASP) and DNA damage markers and were strongly associated with age. p16KB cell percentages were also increased in CD24+ osteolineage cells, which exhibited an inflammatory SASP in aged mice and were robustly cleared by both genetic and pharmacologic senolytic therapies. Following isolation, CD24+ skeletal cells exhibited growth arrest, SA-βgal positivity, and impaired osteogenesis *in vitro*. These studies thus provide a new approach using multiplexed protein profiling by CyTOF to define senescent mesenchymal cells *in vivo* and identify a highly inflammatory, senescent CD24+ osteolineage population cleared by senolytics.

## INTRODUCTION

Cellular senescence is a state of proliferative arrest that occurs due to the accumulation of DNA damage and cellular stress^1–4^. This is distinct from quiescence, as senescent cells can acquire a senescence-associated secretory phenotype (SASP), consisting of pro-inflammatory factors that have detrimental effects on cell and tissue function both locally and systemically^5^. Senescent cells accumulate in the bone microenvironment with age^6^, and clearance of senescent cells in old mice alleviates multiple age-related morbidities^7–12^, increases lifespan^13–15^, and preserves bone microarchitecture and strength^16^. Thus, senescent cell clearance represents a new therapeutic approach to delay or alleviate age-related diseases.

The direct identification of cells undergoing senescence *in vivo* would allow for biological age phenotyping, determination of senolytic efficacy, and identification of novel senolytic targets; however, multiple technical obstacles prevent this process. The variable abundance and rarity of tissue-resident senescent cells^17^ requires large sample sizes to provide sufficient signal and statistical power^18^. Moreover, established markers for senescence (e.g., p16^Ink4a^, p21^cip1^) are expressed intracellularly and at low levels, typically restricting senescence phenotyping to whole-tissue RNA analyses. This method of bulk phenotyping unfortunately lacks the resolution to determine if these cell-cycle proteins are upregulated in the same cells exhibiting a SASP and growth arrest, blurring the distinction between senescence and systemic inflammation. Compounded by their expected heterogeneity^19^, it remains difficult to comprehensively study senescent cells *in vivo* without increased cellular resolution.

Single-cell RNA sequencing technologies have made significant technological advances, yet studies using this approach to investigate senescence present their own challenges. Though overcome by meticulous PCR primer design, the genes encoding both p16^Ink4a^ (*Cdkn2a*) and p21^cip1^ (*Cdkn1a*) proteins generate multiple transcript variants that are challenging to segregate through standard single-cell transcriptomic libraries; *Cdkn2a* also encodes p19^Arf^ (p14^Arf^ in humans), a protein with separate functions and opposing effects on senescence to p16^Ink4a 20–22^, and *Cdkn1a* has multiple variants that associate with aging to varying degrees^23^. Moreover, discrepancies between mRNA and protein expression have been observed for key senescence markers, including p16^Ink4a 24^, which is compounded by the effects of senescence on translation^25^ and proteostasis^26^. Therefore, single-cell techniques leveraging multiplexed protein profiling to assess p16 or p21 proteins, as well as other properties of senescent cells (e.g., the SASP, growth arrest, apoptosis resistance), would potentially be a significant advance for studying the fundamental biology of senescence.

In the present study we leverage the power of mass cytometry by time-of-flight (CyTOF)^27^ to define, at the single-cell level, mesenchymal senescent cells *in vivo*. Specifically, rather than relying principally on p16 or p21 expression, we used multiple criteria to define senescent cells. Moreover, given the inherent biological variability of senescent cells with aging ^28^, we used a large sample size of mice across all experiments (n=88), assessed senescent cell burden in established skeletal cell populations, and tested their susceptibility to senolytic clearance through either genetic or pharmacologic approaches. Findings by CyTOF were supported by single cell RNA-sequencing (scRNA-seq) and *in vitro* phenotyping. Collectively, our studies establish a robust approach to define senescent mesenchymal cells *in vivo* using CyTOF, provide a detailed map of aging- and senolytic-induced alterations in mesenchymal cells in the murine bone microenvironment, and identify specific osteolineage populations that are highly inflammatory, senescent using rigorous criteria, and cleared by senolytics.

## RESULTS

### Development and validation of a senescence CyTOF antibody panel

We constructed and validated a comprehensive CyTOF antibody panel to include markers for both cell identity and senescent phenotype (Table 1). A defining characteristic of senescent cells is expression of cell cycle inhibitors, in particular p16 or p21^29^, so we carefully validated antibodies to these proteins for use in CyTOF. Due to concerns regarding the specificity of antibodies to mouse p16, we tested 3 separate commercial antibodies using an *in vitro* workflow (Fig. 1A-C). Mouse p16 was expressed in human U2OS cells through vector transfection alongside an empty vector control. While some antibodies demonstrated high background and limited positive signal by CyTOF, one antibody in particular provided an excellent signal-to-noise ratio (Fig. 1C) and was therefore selected for the panel. Notably, this antibody has been used in other senescence studies for flow or mass cytometry^30,31^. A similar process was followed for antibodies targeting several additional antigens, including p21 (Extended Data Fig. 1A-D).

**Table 1:**
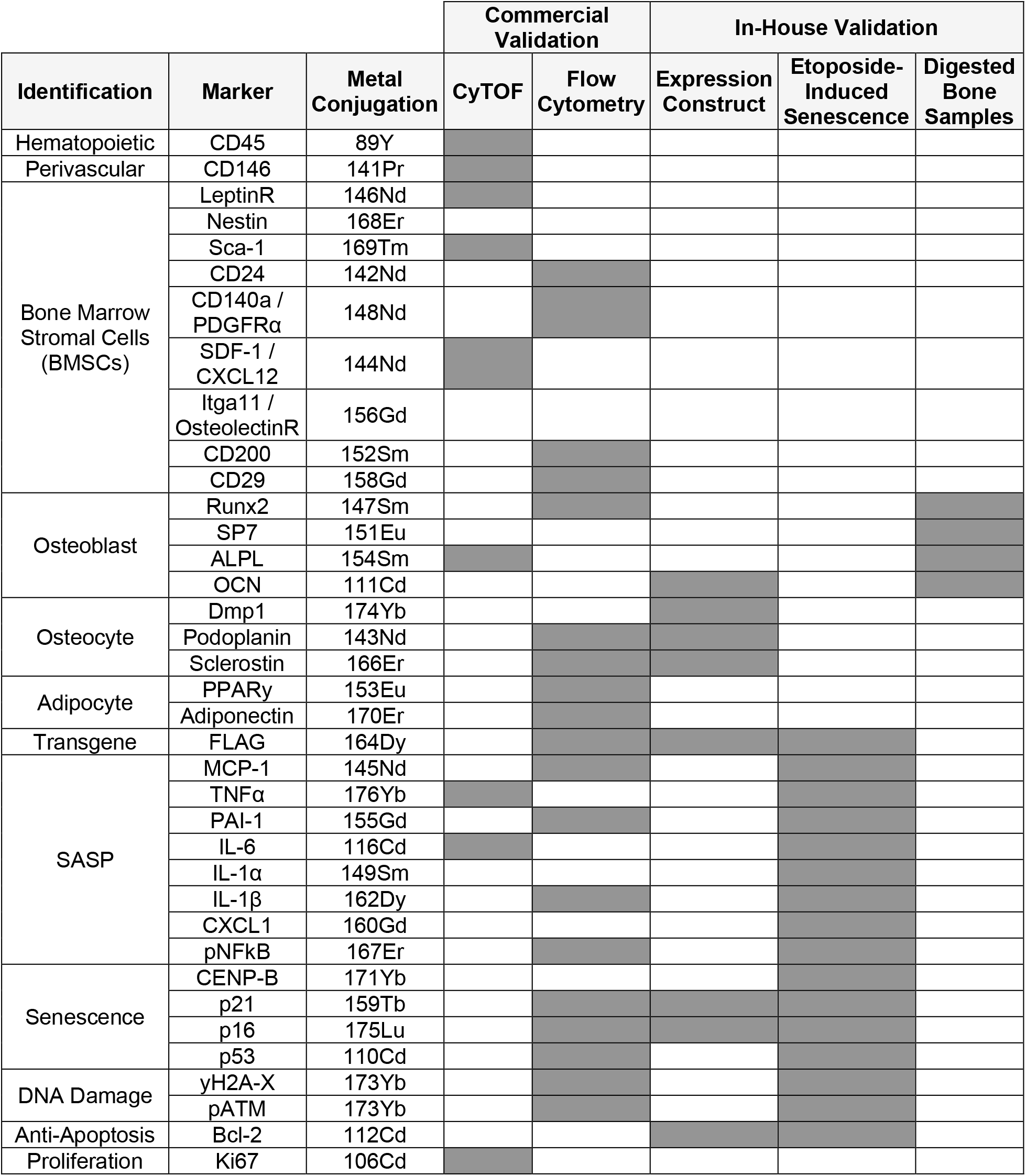
CyTOF Antibody Panel and Validations. Antibodies used in the CyTOF panel were commercially validated for CyTOF/flow cytometry and/or validated in-house using several different approaches.

**Figure 1.**
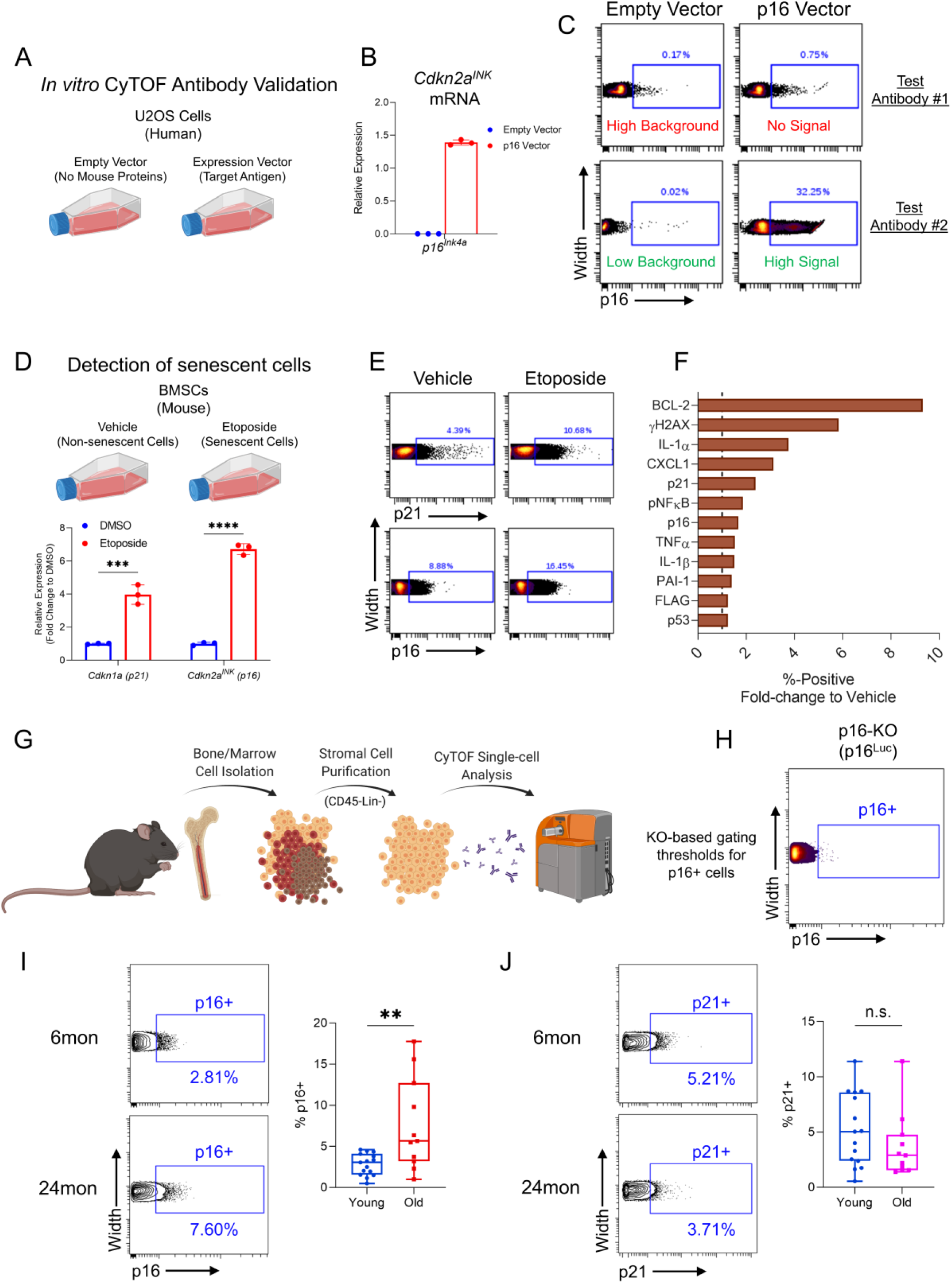
Validation of antibodies and detection of senescent cells by CyTOF. (A) Experimental workflow of single mouse protein expression in U2OS cells for the testing of CyTOF antibodies; (B) qPCR analysis confirming upregulated mouse *p16^Ink4a^* mRNA (*Cdkn2a*) after expression vector transfection; (C) CyTOF dot plots of expression samples testing different p16 antibodies, demonstrating outcomes of both failed (antibody #1) and successful (antibody #2) tests; (D) Schematic of testing senescence-specific CyTOF antibodies using etoposide-treated mouse BMSCs with qPCR confirmation of upregulated *p16^Ink4a^ (Cdkn2a^INK^*) and *p21^Cip1^ (Cdkn1a*) transcripts; (E) CyTOF plots of p16 and p21 protein expression, demonstrating increased percent-positive cells with etoposide treatment; (F) Fold-change of percent-positive values for each of the senescence panel markers in etoposide-treated cells, where dotted line represents values from vehicle. (G) Schematic of bone and marrow mesenchymal cell isolation and CyTOF workflow; (H) Gating strategy for p16+ cells using similarly processed cells from p16-null mice as a negative control (I) Quantification of %p16+ and (J) %p21+ cells in young and old mice; ***p<0.001, ****p<0.0001; (D) Multiple t tests with Holm-Sidak Correction. (I, J) Mann-Whitney test.

To validate our senescence/SASP antibodies and confirm our technical ability to process, stain, and detect senescent cells, our panel was tested on etoposide-induced senescent mouse bone marrow stromal cells (BMSCs) (Fig. 1D). Compared to non-senescent BMSCs, senescent BMSCs demonstrated an upregulation of cells positive for p16, p21, and a majority of our senescence/SASP markers by CyTOF (Fig. 1E, F). We also optimized our single-cell suspension collection protocol from mouse bone and marrow: as shown by others^32,33^, marrow digestion was optimized to release high yields of non-hematopoietic cells from marrow stroma (Extended Data Fig. 2A, B). Additionally, bone tissue digestion was optimized to ensure release of both early (Runx2+, Osterix+) and late (ALPL+, Osteocalcin+) osteolineage cells (Extended Data Fig. 2C-F), which also assisted in validating our osteoblast lineage cell antibodies^34,35^.

We next applied our CyTOF antibody panel to freshly isolated bone and marrow mesenchymal cells from *INK-ATTAC* mice: a transgenic model with an inducible caspase 8 cassette driven by the *p16^Ink4a^* promoter, allowing for selective clearance of senescent cells through treatment with AP20187 (AP)^12^. Using this approach, we could robustly compare cellular alterations resulting from both aging and senolytic clearance within the same mouse strain; accordingly, we collected a total of 40 mice in these groups (n=15 “Young” [6-month], n=12 “Old” [24-month] + vehicle, and n=13 Old [24-month] + AP). We applied our antibody panel to single-cell suspensions of non-immune (Lin- [CD5, CD45R (B220), CD11b, Gr-1 (Ly-6G/C), Ly6B.2 and Ter-119]) and non-hematopoietic (CD45-) stromal skeletal cells (Fig. 1G, see Extended Data Fig. 2G for gating strategy).

Using identically processed cells from p16 knock-out mice^36^ as a negative control (Fig. 1H), we identified p16+ mesenchymal cells from both young and old mice (Fig. 1I). We emphasize that our identification of p16+ cells *in vivo* by CyTOF utilized an antibody that specifically detected the mouse p16 protein in human cells expressing the *p16^Ink4a^* construct (Fig. 1C), demonstrated an increased signal for p16+ cells *in vitro* following etoposide treatment (Fig. 1E), and was thresholded on p16 knock-out cells (Fig. 1H). It is also important to note that the validation of this antibody at this point is only for single-cell CyTOF and other uses (e.g., immunohistochemistry, etc.) would require independent validation of this antibody for that specific purpose. Using this antibody, we found that p16+ cells were more abundant with age, expanding from 2.81% ± 1.32% to 7.60% ± 5.60% (P = 0.004) of the total cell population from 6 to 24 months of age, respectively (Fig. 1I). In contrast, the prevalence of p21+ mesenchymal cells did not increase with age in the bone microenvironment (Fig. 1J; 5.21% ± 3.27% in young, 3.71% ± 2.96% in old [P = 0.241]).

### Single-cell specification of senescent mesenchymal cells in bone and marrow

Although found to be associated with age, p16 positivity can also be a characteristic of non-senescent cells^37–40^, and thus in isolation may not identify a pure population of truly senescent cells. Therefore, to define additional markers uniquely associated with age-induced senescent cells, we performed multidimensional clustering of p16+ mesenchymal cells from young and aged mice using all markers in our senescence panel (Fig. 2A, Extended Data Fig. 3A). We identified considerable cellular heterogeneity, with p16+ cells segregating into 6 unique clusters defined by expression of BCL-2, Ki67, yH2A-X, or other inflammatory markers, along with an unlabeled cluster (Fig. 2B). Across aging, the BCL-2+ and BCL-2+/γH2A-X+ populations increased in prevalence, while the Ki67+ populations were reduced (Fig. 2C). Similar to recent findings using highly sensitive *p16^Ink4a^*-reporter mice (INKBRITE)^37^, we found an inverse relationship between Ki67 and p16 expression (Extended Data Fig. 3B), which also revealed that clusters both high in p16 and low in Ki67 were defined by expression of BCL-2, a major regulator of apoptosis resistance^41^ (Fig. 2D). Fig. 2D also clearly demonstrates that as p16 expression increased across the clusters (red line), Ki67 expression concurrently decreased (blue line). We then performed manual gating on our samples and found that the BCL2+ subset of p16+ cells was inherently reduced in both Ki67 expression and Ki67+ cells (Fig. 2E, F), with nearly 98% of p16+BCL2+ cells being Ki67- (Extended Data Fig. 3C). Across aging, we found that BCL-2 was the only factor clearly upregulated in old versus young p16+ cells (1.8-fold, adj. p=0.0012; Fig. 2G); this finding is entirely consistent with extensive literature demonstrating upregulation of BCL-2 anti-apoptosis pathways in senescent cells^42–44^.

**Figure 2:**
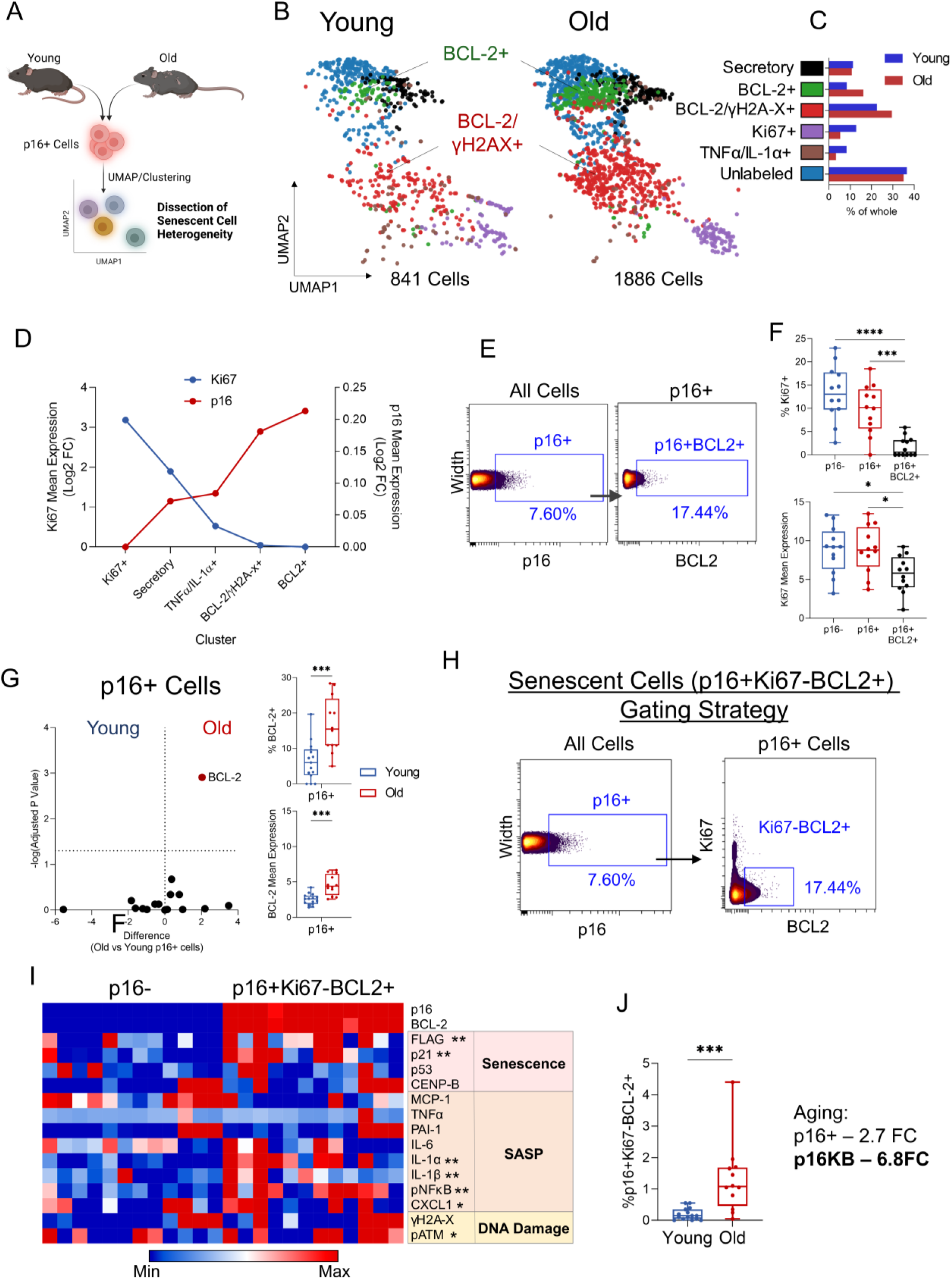
BCL-2 expression defines p16+ cells with senescent characteristics. (A) Schematic of multidimensional p16+ senescent cell analysis workflow; (B) UMAP visualization and FlowSOM clustering of p16+ cells from young and old mice with (C) bar graphs indicating percent-of-whole cluster abundance changes. (D) FlowSOM clusters ranked by descending Ki67 mean expression plotted against p16 mean expression (Log2 Fold-Change). (E) Gating strategy for p16+BCL2+ cells and (F) quantification of %Ki67+ and Ki67 mean expression alongside p16- and total p16+ cells. (G) Volcano plots of age-related changes in mean senescence marker expression within p16+ cells and quantification of % BCL-2+ cells and mean BCL-2 expression within all p16+ cells with age; (H) Gating strategy for “p16KB” cells; (I) Heatmap representation of protein expression between p16KB cells and non-senescent p16-cells, with asterisks indicating significance. (J) Quantification of p16KB cells with age and numerical comparison to total p16+ cells. *p<0.05, **p<0.01, ***p<0.001, ****p<0.0001. (F, G) Multiple t tests with Holm-Sidak Correction. (G, I, J) Mann-Whitney test.

Based on this data, we defined a mesenchymal senescent cell population at the single cell level as being <u>p16</u>+/<u>K</u>i67-/<u>B</u>CL-2+ (hereafter referred to as “p16KB” cells) (Fig. 2H). To support this, we found that p16KB cells exhibited upregulated expression of numerous markers for senescence (p21), SASP (IL-1α, IL1β, pNFκB, CXCL1) and DNA damage (pATM) (Fig. 2I). Moreover, p16KB bone and marrow cells were highly age-associated, making up only <0.2% of all cells in young mice, yet with a fold-change of 6.8 across aging (Fig. 2J) (as compared to total p16+ cells: 2.81% in young, 2.7-fold with age, Fig. 1I).

Similar to p16, BCL-2+ subsetting of p21+ cells identified a population with reduced Ki67 positivity compared to total p21+ cells (Extended Data Fig. 3C). Analogous p21+/Ki67-/BCL-2+ (“p21KB cells”) were identified (Extended Data Fig. 3D), which displayed a robust SASP and DNA damage expression profile (Extended Data Fig. 3E). Interestingly, despite no age-related increase in total p21+ cells, p21KB cells did increase significantly in old mice (Extended Data Fig. 3F), thus identifying an age-associated p21+ subpopulation. However, p16KB cells greatly outnumbered p21KB cells in old mice and exhibited a more dramatic upregulation with age (Extended Data Fig 3G). Upon further investigation, we found minimal overlap between p16KB and p21KB cells (0.04% of total cells in young, 0.10% in old; Extended Data Fig. 3H). To test the validity of both p16KB and p21KB results, we replicated this CyTOF analysis in an additional cohort of young (6-month) and aged (24-month) C57BL/6N mice (wild-type, rather than *INK-ATTAC*) (Extended Data Fig. 4A). We found our observations to be robustly conserved, revealing similar age-associated patterns for total p16+ and p21+ cells, with BCL-2 subsetting identifying growth-arrested and age-associated subpopulations for each (Extended Data Fig. 4B-D).

Collectively, using single cell mass cytometry, we thus define p16KB and p21KB mesenchymal cells as being senescent, as these populations fulfill virtually all of the required criteria for senescent cells^45^: increased p16 or p21 protein expression, growth arrest, upregulation of anti-apoptotic pathways, upregulation of SASP and DNA damage markers, and near absence in youth with a marked increase with aging. Moreover, we note that in addition to our use of carefully validated antibodies in the CyTOF analysis, our orthogonal approach to identify p16KB and p21KB cells does not rely solely on the specificity of any single antibody, but rather uses a combinatorial approach, making it extremely unlikely that the p16KB or p21KB senescent cells represent a false positive signal.

### Reconstruction of bone and marrow mesenchymal populations through CyTOF

Next, in order to assess which mesenchymal skeletal cell populations harbored senescent cells with age, we established our bone-resident clusters. Using t-SNE visualization and FlowSOM cell clustering, we identified 11 populations of BMSCs (also termed skeletal stem cells [SSCs]) and differentiated cell types with expression profiles consistent with the established literature (Fig. 3A, B; see Table 2 for a detailed description of defining markers for each population). These included LeptinR+ BMSCs^46^, Sca-1+/PDGFRα+ BMSCs (also termed PαS cells)^47–49^, perivascular BMSCs^50,51^, and Nestin+ pericytes^52^. Other characteristics of these cells have been established by recent studies, such as expression of CXCL12^53^ and OsteolectinR (*Itga11*)^54^ in LeptinR+ BMSCs, Adiponectin expression in perivascular and stromal cells^55^, and PDPN expression in Sca-1/PDGFRα+ BMSCs^49,56^. Additional clusters representing committed cell types included early osteoblasts (Runx2+/Osterix+)^57,58^, alkaline phosphatase (ALPL)+ osteolineage cells (Runx2+/ALPL+)^59,60^, late osteoblasts/osteocytes (Runx2+/Osterix+/Sclerostin+; note that our cell isolation protocol included a collagenase digestion thereby releasing at least a subset of osteocytic cells)^61^ and pre-adipocytes (Pparg+)^62,63^. Other populations represented less well-defined populations, including CD24^high/low^ osteolineage (CD24+/Runx2+/Osterix+) and CD24+/Osterix+ clusters. CD24 has previously been linked to osteogenesis^64^, while conversely also shown to mark pluripotent BMSCs^65,66^, particularly when co-expressed with Sca-1^67^ or CD200; however, these CD24 clusters lacked co-expression of these stem-like markers (Extended Data Fig. 5A, B) indicating a distinct, more committed osteoblast lineage population.

**Figure 3:**
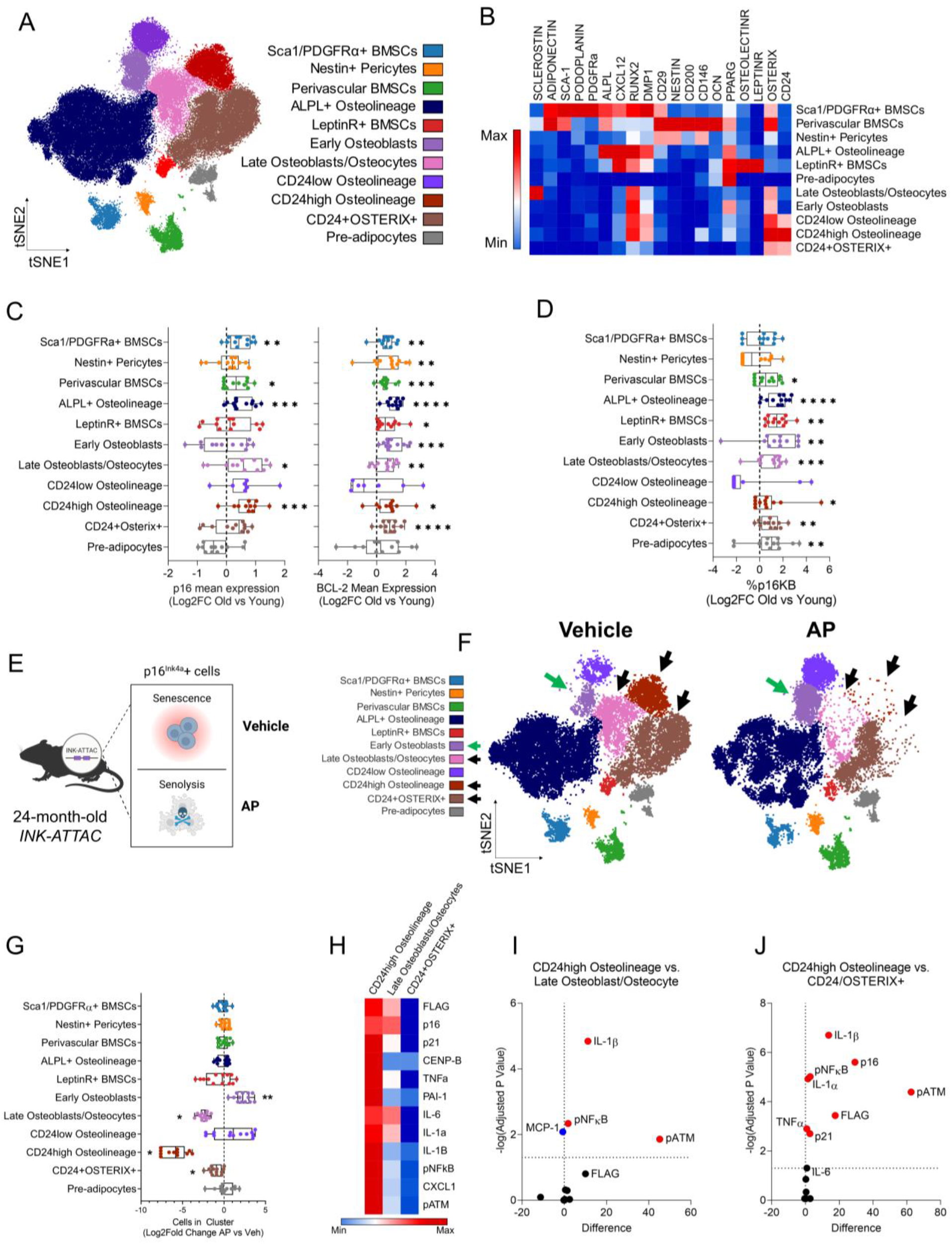
CD24^high^ osteolineage cells represent inflammatory senescent cells in old mice targeted by genetic senolytic clearance. (A) t-SNE visualization and FlowSOM clustering of n=80,000 CD45-Lin-bone and marrow cells (n=40 INK-ATTAC mice [n=15 young, n=12 old + vehicle, n=13 old + AP] – 2,000 cells sampled per mouse) analyzed by CyTOF. Cells are colored by clustered population (see Table 1 for defining markers); (B) Heatmap representation of the 11 cell clusters and protein expression of identification markers; (C) Log2 fold-Change mean expression of p16 or BCL-2 in each cluster with age. (D) Log2 fold-change of %p16KB cells in each cluster across aging; (E) Schematic of p16+ senescent cell clearance in 24-month-old AP-treated *INK-ATTAC* mice; (F) t-SNE plots of FlowSOM clusters of bone/bone marrow cells from old vehicle- or AP-treated *INK-ATTAC* mice. Black arrows indicate cleared clusters while green arrows indicate increased cluster abundance; (G) Quantification of cluster abundance changes between vehicle- and AP-treated mice (Log2-fold change to vehicle); (H) Heatmap and (I, J) volcano plots for statistical comparisons of senescence marker expression between cleared clusters in vehicle-treated old mice. *p<0.05, **p<0.01, ***p<0.001, ****p<0.0001; (C, D) Mann-Whitney test or Unpaired t test, as appropriate. (G, I, J) Multiple t tests with Holm-Sidak Correction.

**Table 2:**
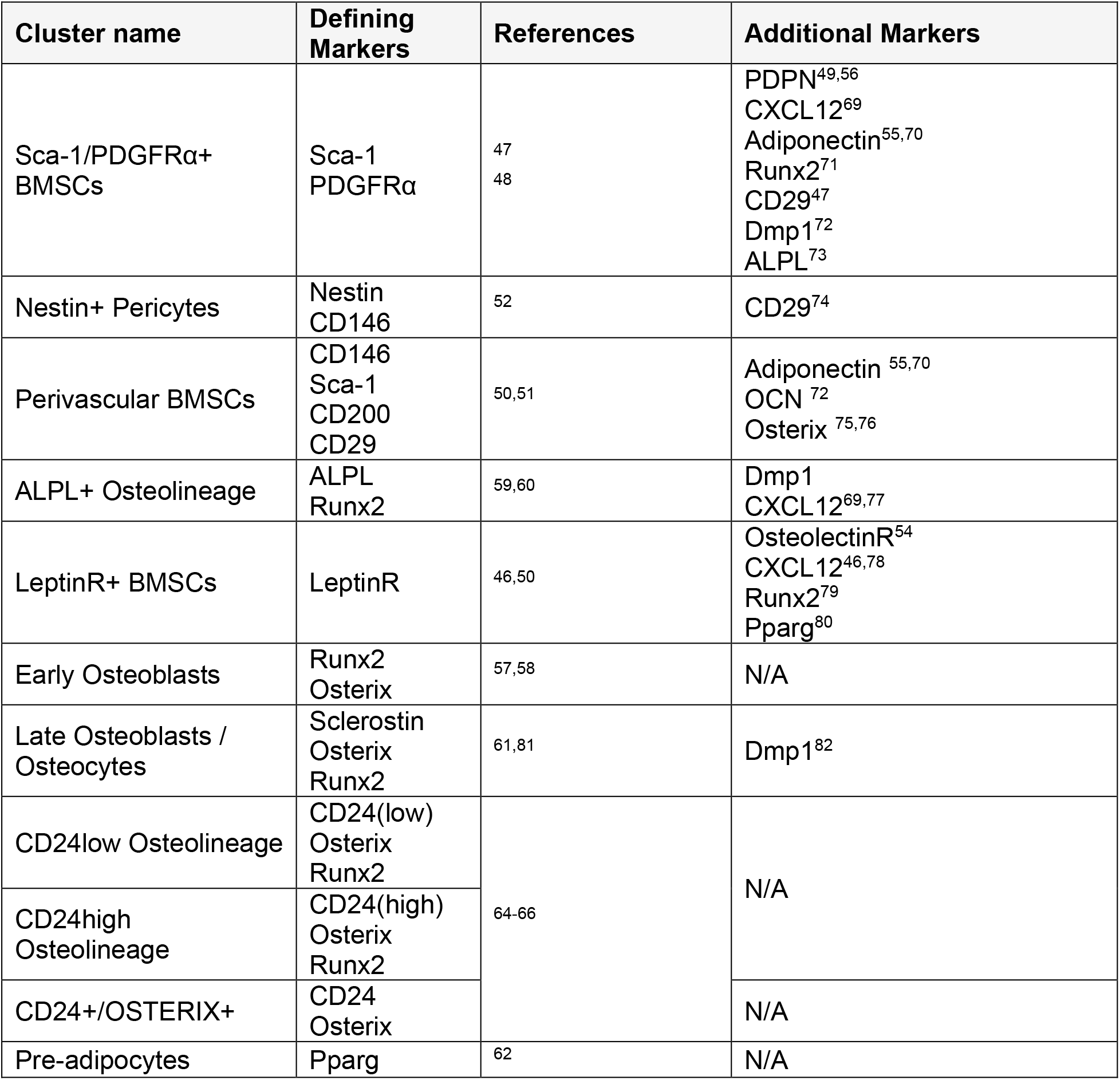
Defining markers for CyTOF cluster identification. Each cluster was defined by expression of specific defining markers and additional markers as characterized by previous studies in the literature.

We next evaluated our CyTOF panel for its ability to delineate the various stages of mesenchymal cell differentiation. Diffusion mapping displayed perivascular BMSCs and Nestin+ pericytes on one end of the lineage continuum, and Sca-1/PDGFRα+ BMSCs on another, converging to form several committed osteogenic clusters (Extended Data Fig. 5C). This is consistent with many studies that demonstrate the multipotency of these populations and indicate multiple sources of osteolineage cells^68^. To determine directionality, we performed pseudotime analysis and trajectory inference, seeking to recapitulate the dynamics of *in vivo* cell differentiation. The progression of our clusters was consistent with our current understanding of biological mesenchymal differentiation of skeletal stem cells (Extended Data Fig. 5D, E): BMSC clusters are present early, then leading to early osteoblasts, ALPL+ osteolineage, and ultimately late osteoblast/osteocyte clusters. Interestingly, it appeared that the CD24+ clusters formed their own bifurcation from the typical differentiation of BMSCs to late osteoblasts and osteocytes (Extended Data Fig. 5D) and the CD24+ populations appeared to be middle (CD24^low/high^ osteolineage [CD24+/Runx2+/Osterix+]) to later (CD24+/Osterix+) in the pseudotime progression (Extended Data Fig. 5E), consistent with their lack of expression of stem cell markers such as Sca-1 or CD200 (Extended Data Fig. 5A, B).

To further validate our CyTOF clusters, we performed single-cell RNA-sequencing (scRNA-seq) on a similarly prepared sample of bone and marrow cells from aged mice (Extended Data Fig. 6A). After multidimensional analysis, we found that skeletal cell populations identified through single-cell transcriptomics were entirely consistent with our CyTOF clusters (Extended Data Fig. 6B). Specifically, the BMSC populations (Sca-1/Pdgfra+, Perivascular, and Nestin+ Pericytes) and CD24 osteolineage populations (*Cd24a, Runx2*) appeared well conserved, in addition to the distribution of expression for certain markers (e.g. *Cxcl12*) (Extended Data Fig. 6C). However, some markers were expressed at very low/undetectable levels (e.g. *Dmp1, Sp7* [Osterix], *Sost*) preventing classification of late osteogenic clusters. Besides these limitations by scRNA-seq, the overall population structure was comparable to that which we observed using CyTOF, adding confidence that this proteomic approach provides accurate interpretations of the bone microenvironment.

### Mature osteolineage populations harbor age-induced senescent cells

To identify mesenchymal populations affected by aging, we next examined senescence-specific effects, focusing on p16KB senescent cells, as we have previously demonstrated that clearing p16+ cells in *INK-ATTAC* mice prevents age-related bone loss, reduces bone resorption, and increases bone formation^16^. We found that multiple clusters exhibited upregulation of p16 and/or BCL-2 expression with age (Fig. 3C), with many of these clusters exhibiting an expected increase in the percentage of senescent p16KB cells with age (Fig. 3D).

We next tested if the above populations were those that are cleared in aged *INK-ATTAC* mice (Fig. 3E; Extended Data Fig. 7A). Upon treatment with AP, cells in the late osteoblast/osteocyte cluster were markedly reduced (Fig. 3F), which is consistent with our previous data demonstrating a central role of osteocyte senescence with age^6,16,83–85^. In addition to this confirmatory result, we also uncovered that the CD24high Osteolineage population was markedly reduced following AP treatment (Fig. 3F, G), along with a modest reduction in CD24+Osterix+ cells. In addition to clearance of these clusters, the Early Osteoblast cluster increased in number after AP treatment (Fig. 3F, G), suggesting that this is a newly replenished osteogenic population that appears after the clearance of senescent cells. This is supported by pseudotime analyses, which indicated that there is an emergence of cells that are early in the differentiation continuum following clearance of p16+ senescent cells (Extended Data Fig. 7B). This finding is entirely consistent with our previous work which demonstrated that AP treatment in aged *INK-ATTAC* mice improved endosteal osteoblast numbers and bone formation rates^16^. Finally, to support these findings we performed CITRUS analysis, which generates separately stratified clusters from the original dataset to observe statistical differences, and independently found downregulation of CD24+ clusters with AP treatment, along with an increase in Runx2+ and Runx2/Osterix+ early osteoblast clusters (Extended Data Fig. 7C).

Importantly, as noted above, we have previously demonstrated senescence within osteocytes with age – and their clearance with senolytic therapy – through gene expression and histological senescent phenotyping^16^. Entirely consistent with this, we now demonstrate concomitant increases in p16 and BCL-2 expression (Fig 3C) and in %p16KB cells (Fig 3D) by the late osteoblast/osteocyte cells, along with clearance of these cells following AP20187 treatment (Fig 3E-G). In addition, we identify new populations (CD24high osteolineage /CD24+Osterix+ cells) that also demonstrate increases in %p16KB cells with age (Fig 3C, D) and are robust targets of senolytic clearance (Fig 3E-G). Interestingly, though, in aged mice, CD24^high^ osteolineage cells displayed the highest expression levels of markers for senescence, SASP, and DNA damage, with the CD24+Osterix+ cells being the lowest, and the late osteoblast/osteolineage cells having intermediate levels of these markers (Fig. 3H-J). These data thus reveal that CD24^high^ osteolineage cells exhibit a robust senescent and inflammatory profile relative to other skeletal populations and - in addition to osteocytes, as previously reported^16^ - these CD24^high^ osteolineage cells are key targets of p16-driven senolytic clearance.

### CD24 osteolineage populations exhibit a unique SASP profile with age

To further investigate the senescent profile of these CD24+ osteolineage cells, we applied CITRUS analysis and compared this to our established FlowSOM clusters (Fig. 4A). Consistent with the findings noted above, we found that many clusters exhibited higher expression levels of senescence/SASP proteins in old compared to young mice (FDR<5%) (Extended Data Fig. 7D). Of these proteins, p16 and BCL-2 were upregulated in the largest number of clusters with age, supporting their use as robust predictors of senescence (Fig. 4B). These factors were followed closely by CXCL1, FLAG (detecting the *p16^Ink4a^* promoter activity in the *INK-ATTAC* transgene) and several interleukins (IL-6, IL-1α, IL1β). As observed previously, p21 was not found to be differentially regulated with age by CITRUS.

**Figure 4.**
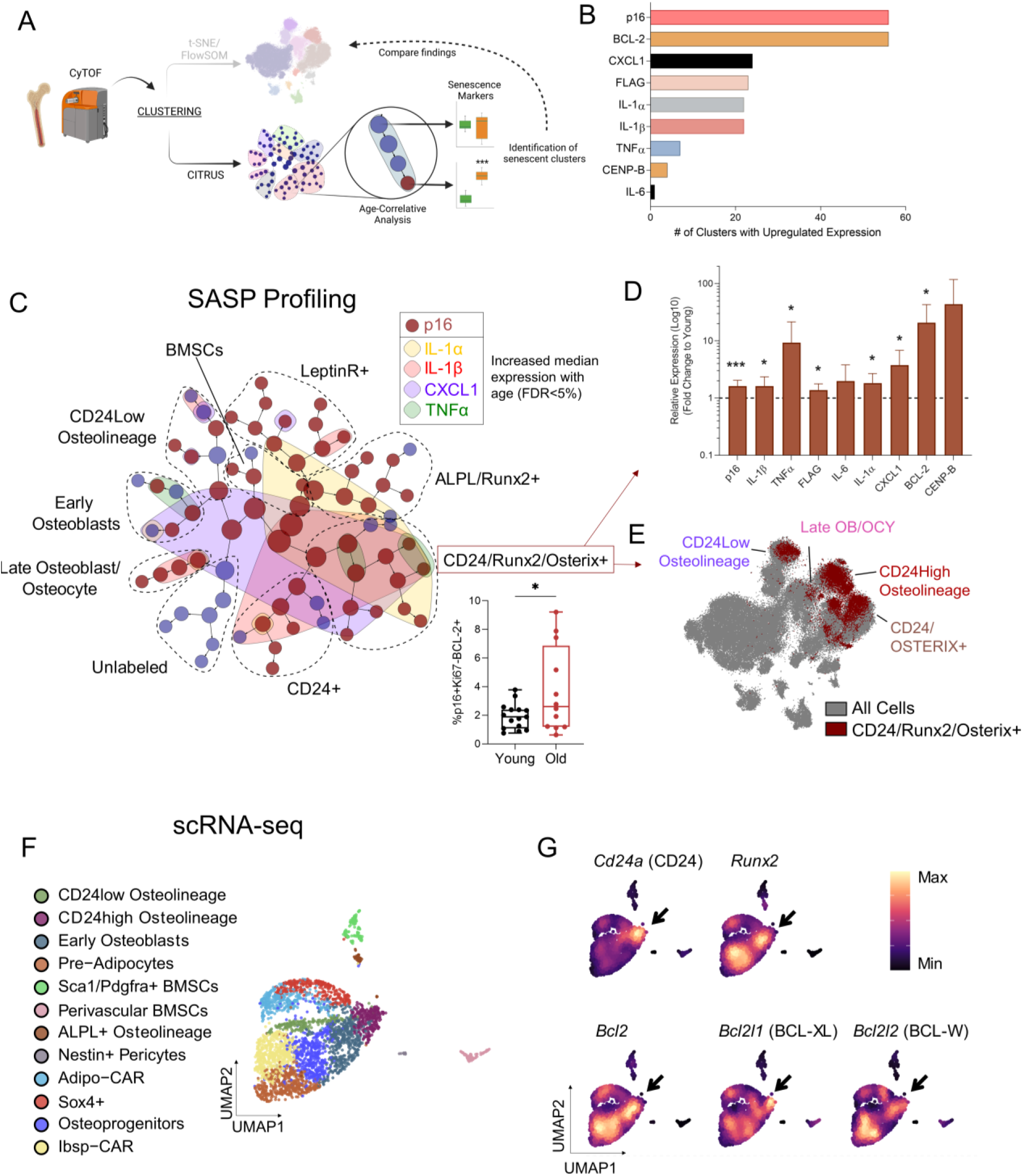
CITRUS and scRNA-seq analyses. (A) Schematic of CITRUS analysis workflow; (B) Aging markers upregulated in the highest number of CITRUS clusters with age (FDR<5%); (C) Senescence/SASP marker expression changes across aging overlaid on CITRUS plot (each circle is a cluster, grouped by cluster families dotted lines; see Figure S6A for identity marker expression). Red clusters indicate upregulated p16 expression, while colored overlays indicate upregulation in the corresponding SASP marker. All markers converge in CD24+/Runx2+/Osterix+ clusters, indicating that this population exhibits the most widespread age-related increase in SASP marker expression in bone/bone marrow. Box plot below demonstrates an upregulation of %p16KB cells with age in this population. Red arrows point to (D) quantified median expression of various SASP factors in the CD24+/Runx2+/Osterix+ clusters with age and (E) overlay of the CD24+/Runx2+/Osterix+ cluster on original t-SNE plot with FlowSOM clusters labeled; (F) UMAP visualization by scRNA-seq of n=3,362 clustered Lin-CD45-cells from the digested bone and marrow of n=3 24-month untreated *INK-ATTAC* mice; CAR, Cxcl12-abundant reticular; (G) Density plots of CD24 osteolineage markers and anti-apoptotic factors (*Bcl2, Bcl2l1, Bcl2l2*). *p<0.05, ***p<0.001. (C) Mann-Whitney test. (D) Multiple t tests with Holm-Sidak Correction.

Among the clusters with increased p16 expression with age, cells expressing CD24/Runx2/Osterix exhibited a unique upregulation of the most SASP inflammatory proteins, including IL-1α, IL1β, CXCL1, and TNFα (Fig. 5C, D); these cells also exhibited robust upregulation of BCL-2 and an increased percentage of p16KB cells with age (Fig. 4D). When these CD24/Runx2/Osterix cells were overlaid with our original t-SNE plots, they were largely contained within CD24+ osteolineage clusters, along with partial overlay in the late osteoblast/osteocyte cluster (Fig. 4E). This independent CITRUS analysis thus strongly supports our previous data demonstrating increased senescent cell burden in these clusters with age (Fig. 3C, D) and their clearance with AP (Fig. 3F).

**Figure 5.**
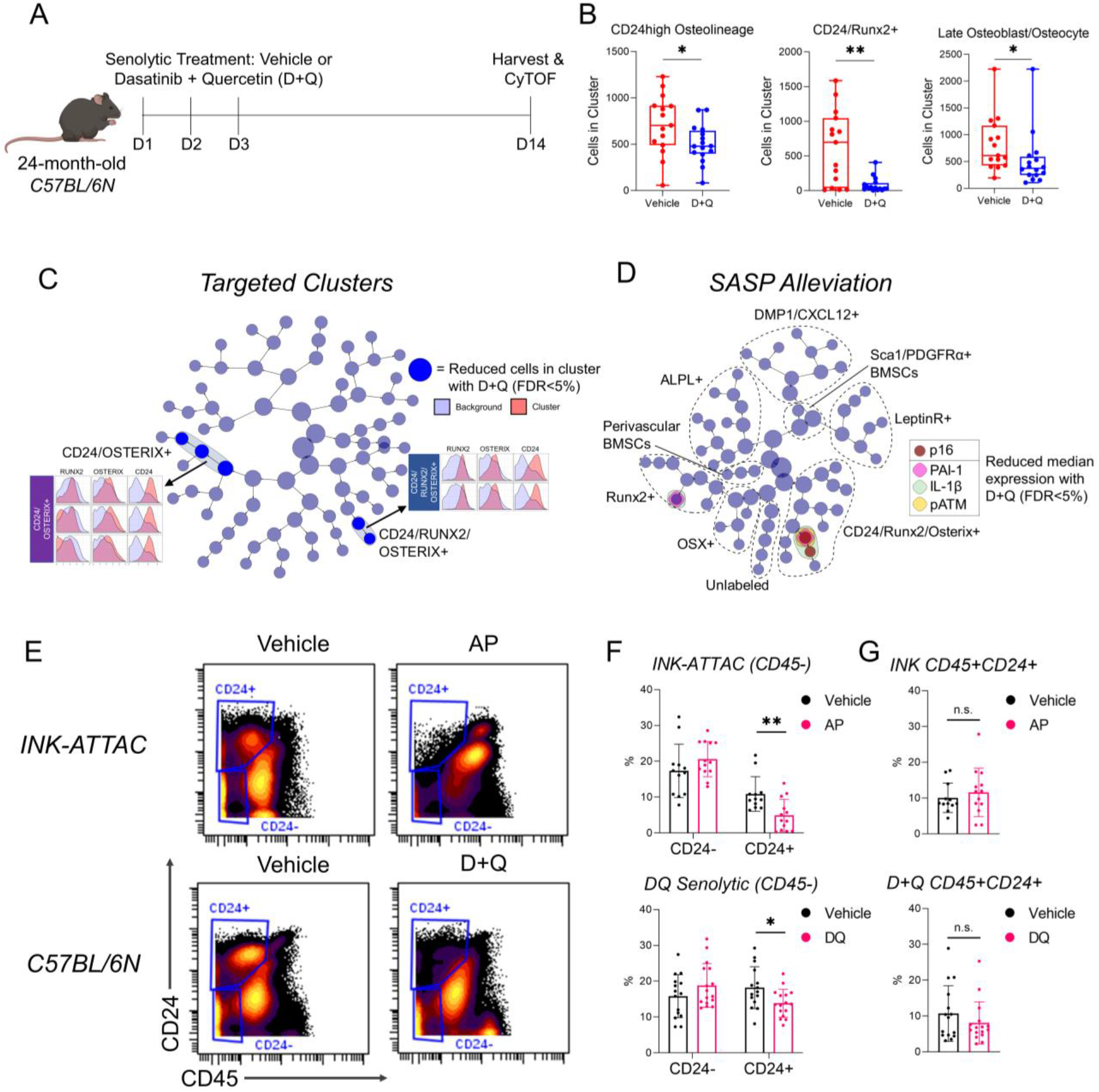
Pharmacologic senolytic therapy targets CD24+ osteolineage cells in aged mice. (A) Experimental design of pharmacological senolytic treatment of 24-month-old C57BL/6N mice with Dasatinib + Quercetin (D+Q). Mice were treated for 3 consecutive days, then harvested at 14 days (indicated by dashes); (B) Quantified cell abundance changes with D+Q treatment in clusters similarly targeted in *INK-ATTAC* mice (See Extended Figure 8A, B for cluster definitions); (C, D) CITRUS analysis of cluster abundance (C) and median expression changes (D) between vehicle- and D+Q-treated mice (FDR<5%). (C) Cleared clusters, marked by blue, are defined by high CD24, Osterix, and/or Runx2 expression, as shown by histograms. (D) Clusters with reduced median expression are colored by their respective marker. Cluster families are marked by dotted lines (See Extended Figure 8C for defining markers); (E) Gating strategy for CD45-CD24+ and – populations from all cells in CyTOF data from *INK-ATTAC* mice treated with vehicle or AP, and C57BL/6N mice treated with vehicle or D+Q; (F) Quantification of cell population percentages demonstrate both senolytic treatments clear CD45-CD24+ cells, but not CD45-CD24- or (G) CD45+CD24+ cells. *p<0.05, **p<0.01; (B, G) Mann-Whitney or Unpaired t test, as appropriate; (F) Multiple t tests with Holm-Sidak Correction.

We next investigated our scRNA-seq dataset, finding that CD24 osteolineage cells (positive for *Cd24a* [CD24] and *Runx2*) strongly co-expressed not only *Bcl2*, but also other anti-apoptotic genes *Bcl2l1* (BCL-XL) and *Bcl2l2* (BCL-W) (Fig.4F, G). This suggests that in addition to their inflammatory profile, CD24 osteolineage cells are strongly resistant to apoptosis, perhaps through several mechanisms. Collectively, we further establish that CD24 defines a senescent osteolineage population exhibiting an expression profile enriched for SASP and apoptosis-resistance factors in the aging bone microenvironment.

### Pharmacological senolytic treatment also targets CD24^high^ Osteolineage cells

The unique inflammatory profile of senescent CD24 osteolineage cells led us to test if they are targeted not only by genetic but also by pharmacological senolytic treatment, which selectively kills cells based on senescent cell anti-apoptotic pathways (SCAPs) rather than by activation of transgenic caspase 8 in p16+ cells (as done in *INK-ATTAC* mice). Thus, we performed CyTOF on aged *C57BL/6N* wild type mice treated with or without Dasatinib + Quercetin (D+Q) (Fig 5A): a combination senolytic therapy that targets SCAPs and which we have previously shown to reduce frailty and prevent bone loss in aged mice^14,16,86^. Multidimensional analysis from this cohort identified skeletal cell populations consistent with our *INK-ATTAC* cohort (Extended Data Fig. 8A, B), and treatment with D+Q similarly targeted CD24^high^ Osteolineage, CD24+/Runx2+, and late osteoblast/osteocyte clusters (Fig. 5B). Independently, with CITRUS analysis we found that D+Q was more limited than the genetic *INK-ATTAC* model in its clearance, only targeting 5 clusters overall (Fig. 5C). These clusters were all high in expression for CD24 and represented two separate families of CD24+/Osterix+/Runx2+ and CD24+/Osterix+ clusters. In sequential analyses, we found that D+Q reduced expression of p16, SASP factors PAI-1 and IL-1β, and the DNA damage marker pATM within CD24/Runx2/Osterix+ cells (Fig. 5D; Extended Data Fig. 8C). This aligns with the established effectiveness of D+Q on senescent cells exhibiting DNA damage and serpine (PAI) family proteins^86^.

Due to the recurring presence of CD24 on senescent cell clusters, we next sought to determine if this marker can be used to enrich for cells susceptible to senolytic clearance. Using manual gating on our CyTOF data, we found that total CD45-CD24+ stromal cells were reduced in both genetic (*INK-ATTAC*) and pharmacological (D+Q) methods of senolytic clearance, while CD24– cells were unaffected (Fig. 5E, F). Importantly, CD24+ cells that were CD45+ were not cleared (Fig. 5G), demonstrating senolytic specificity for non-hematopoietic, mesenchymal CD24+ cells. Overall, we establish that CD24 is a consistent marker on aged skeletal senescent cells that are cleared by both genetic and pharmacologic senolytic therapy.

### CD24+ Cells Display Functional Characteristics of Senescence

To further test if CD24+ skeletal cells are enriched for growth arrested senescent cells, we isolated non-hematopoietic (CD45-/Lin-) CD24+ and CD24-cells from the digested bone and marrow of aged mice for *in vitro* phenotyping (Fig. 6A; See Extended Data Fig. 8D for gating strategy). Cell cycle analyses revealed that CD24+ cells were largely growth arrested, with a larger proportion of cells in the G0/1 phase and less in the G2/M phase than CD24-cells (Fig. 6B, C). When placed in culture, CD24+ cells exhibited markedly reduced colony forming efficiency (CFE) after 7 days, while CD24-cells grew rapidly (Fig. 6D-E). CD24+ cells generated very small colonies, typically with only a few cells, demonstrating impaired stemness and proliferative capabilities. Moreover, CD24+ cells exhibited spontaneous senescence, with up to 40% of cells staining positive for senescence-associated β-galactosidase (SA-β-gal) after only 14 days in culture, while CD24-cells continued to proliferate (Fig. 6E).

**Figure 6.**
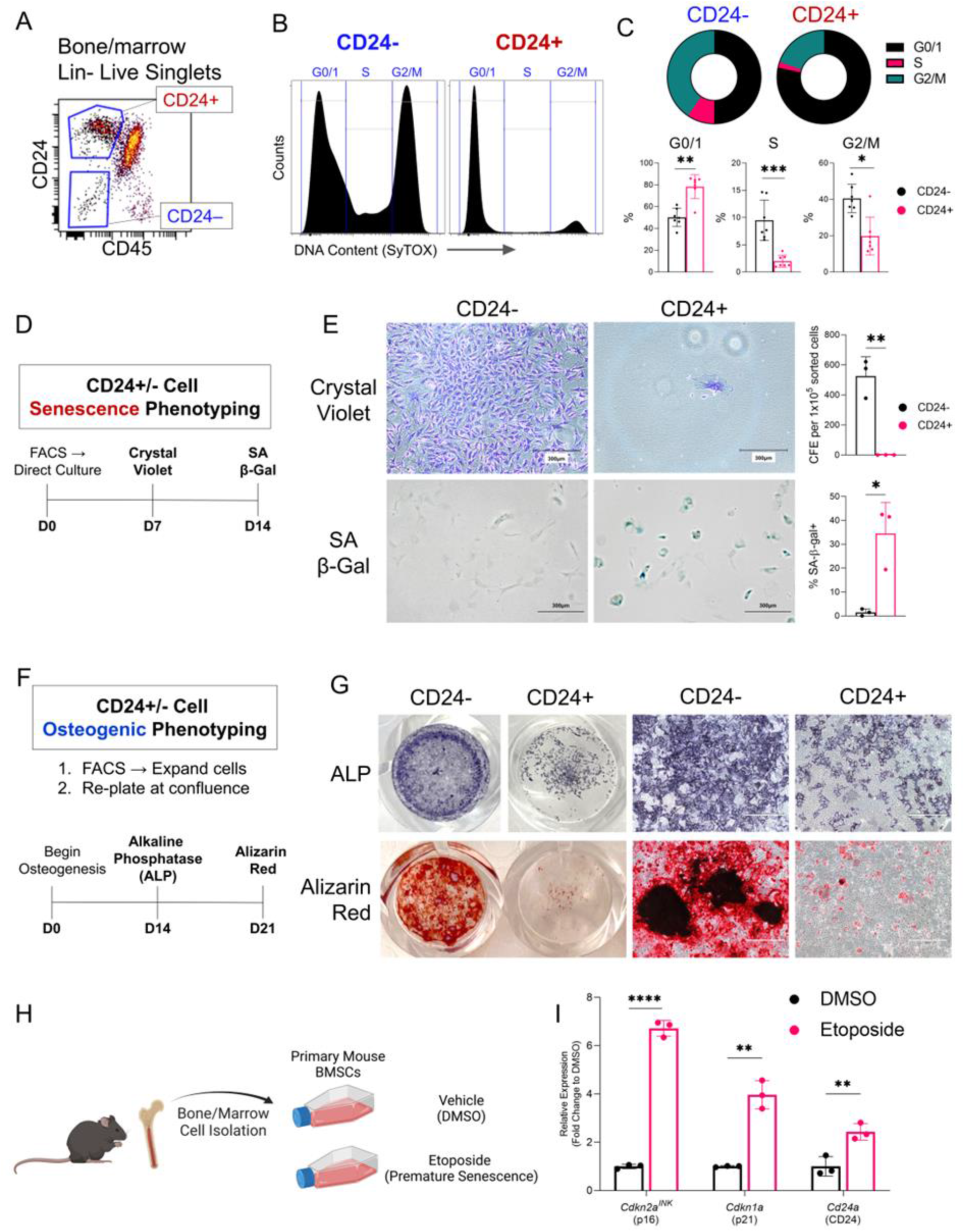
Isolated CD24+ bone stromal cells exhibit functional characteristics of senescence and impaired osteogenesis. (A) Gating strategy for FACS isolation of CD24+ and – skeletal stromal cells from Lin-depleted bone/marrow cell suspensions; (B) Gating strategy and (C) quantification of cell cycle analysis of CD24+/- cells (n=7 mice); (D) Outline of *in vitro* senescence phenotyping of CD24+/- cells; (E) Brightfield images and quantification of colony formation efficiency (CFE) assay and SA-β-gal staining of CD24- and CD24+ cells after 7 or 14 days in culture, respectively (n=3 mice); (F) Outline of osteoblast differentiation assays of CD24+/- cells; (G) CD24- and + cell monolayers stained with either alkaline phosphatase (ALP) or Alizarin Red after 14 and 21 days in culture, respectively. Accompanying high magnification images are 20X (n=3 mice); (H) Schematic of *in vitro* etoposide-induced senescence of BMSCs; (I) mRNA levels of *Cdkn2a^INK^* (p16), *Cdkn1a* (p21), and *Cd24a* (CD24) from BMSCs treated with vehicle (DMSO) or etoposide (n=3 mice). Scale bars represent 750μm. Bars show mean ± SD. *p<0.05, **p<0.01, ***p<0.001, ****p<0.0001. (C, E) Unpaired t test or Mann-Whitney test as appropriate; (I) Multiple t tests with Holm-Sidak Correction.

Although co-expressed with osteogenic markers in both scRNA-seq and CyTOF data, it remains unclear if CD24+ cells have osteogenic capabilities. Thus, we induced osteogenesis in both CD24+ and CD24-cells (Fig. 6F), finding that CD24+ cells have limited osteogenic potential (Fig. 6G). CD24-cells underwent robust osteogenesis, staining strongly for alkaline phosphatase (ALP), a measure of osteoblast differentiation, and Alizarin Red, which stains mineralized nodules. In comparison, CD24+ cells exhibited impaired ALP and Alizarin Red staining, even with sufficient cell numbers (Fig. 6G). Interestingly, these *in vitro* findings are consistent with the *in vivo* pseudotime analysis in Extended Data Fig. 5D which shows that the CD24+ osteolineage cells formed their own bifurcation distinct from the typical differentiation of BMSCs to late osteoblasts and osteocytes.

In various cancers, CD24 serves as a stress marker indicative of tumor burden, rather than a cell-specific marker^87–89^, which led us to test if CD24 may serve a similar role in senescence. We found that CD24 is induced in the senescence program, as *in vitro* etoposide-induced senescence of BMSCs led to elevated *Cd24a* expression, alongside *Cdkn2a^INK^* (p16) and *Cdkn1a* (p21) (Fig. 6H-I). Taken together, these data demonstrate that CD24+ stromal cells exhibit functional characteristics of senescence and impaired osteogenesis, and CD24 may label cells prone to senescence and senolytic clearance due to its upregulation during the onset of senescence.

## DISCUSSION

As senescent cells cannot be defined by a singular marker^90^, the identification of these cells *in vivo* has been impeded by technical limitations. Here, we provide a new approach to identify and characterize *in vivo* senescent mesenchymal cells using multiplexed cellular profiling by CyTOF. Using a methodically validated antibody panel, we dissected senescent cell heterogeneity and defined specific cell populations fulfilling major requirements for senescence. We then applied this definition to aging skeletal cell populations to provide new insights into the identity and characteristics of senescent mesenchymal cells in the bone microenvironment, uncovering physiologically-relevant cell types that are targeted by senolytic treatments we previously demonstrated to be effective in preventing age-related bone loss in mice^16^.

Although long-established as a marker of senescence, we found that p16 expression labels a diverse set of cells with both senescent and non-senescent properties. This finding is entirely consistent with recent findings using highly sensitive *p16^Ink4a^* reporter mice (INKBRITE)^37^, although we provide important complementary data at the protein level to the transcriptional data in the previous study. Specifically, similar to our finding of p16 protein expression in both Ki67 negative and positive bone and marrow mesenchymal cells, in the INKBRITE *p16^Ink4a^* reporter mice, *p16^Ink4a^* was also expressed in both proliferating and non-proliferating lung fibroblasts. Importantly, and entirely consistent with our findings showing an inverse association between p16 and Ki67 protein expression (Fig. 2D-F; Extended Data Fig. 3B), high *p16^Ink4a^*-expressing fibroblasts from the INKBRITE mice had lower proliferative capacity than low *p16^Ink4a^*-expressing cells^37^. Collectively, these findings indicate that there is a spectrum of *p16^Ink4a^* RNA and protein expression *in vivo*, with high expression associated with growth arrest and the full senescent phenotype.

Our finding that BCL-2 co-expression is required to define senescent p16+ cells that are also growth-arrested (Ki67-) aided in our further defining of senescent skeletal cell types, without additional noise through comparing total p16 positivity. Moreover, our strict validation of the p16 antibody for antigen specificity, sensitivity, and biological signal permitted the in-depth characterization of these cells. The application of this CyTOF panel therefore could advance the study of senescence in various murine disease states without the need for genetic reporters.

Like p16+ cells, p21+ cell subsets positive for BCL-2 were associated with age in skeletal mesenchymal cells, even though total p21+ cells were not. Although this was unexpected, as p21 has long been established as a marker for senescence^29,91–93^, this further exemplifies the ability of BCL-2 co-expression to define senescent cells. Furthermore, although this work focused on p16KB cells due to their higher abundance, greater association with age, and previous studies demonstrating beneficial skeletal effects in aged mice of clearing p16+ cells^16^ it will be of interest for future studies to define the function of p21KB cells in skeletal aging. Recent work has demonstrated that p21+ senescent cells are causal to radiation-induced bone loss, while p16+ senescent cells are dispensable^94^. These data suggest the hypothesis that, in bone (and perhaps in other tissues), p16KB cells contribute to age-related senescence, while p21KB cells, albeit still senescent, may have more of a prominent role in acute senescence caused by injury. This is supported by recent work demonstrating that p21 expression is upregulated in bone fracture, with its highest expression immediately following injury, and then waning as p16 upregulation emerges^95^.

Our work also establishes CD24 osteolineage cells as a previously uncharacterized senescent skeletal cell population in aged mice, which we characterized both *in vivo* and *in vitro*. Isolated CD24+ cells exhibited an enrichment of senescent cells and had limited osteogenic capabilities. *In vivo*, we found that the CD24 osteolineage cells had a marked upregulation of SASP factors and clearance of these cells resulted in an increase in early osteoblast lineage cells, consistent with our previously observed effects of senescent cell clearance on improving bone formation in aged mice^16^. Additional *in vitro* and *in vivo* studies are needed, however, to further define the possible paracrine effects of these CD24 osteolineage cells on osteoblasts as well as osteoclasts and other cells in the bone microenvironment, including adipocytes and hematopoietic cells.

CD24 is a cell-surface marker with established roles in stress and immune signaling^87,88,96–99^, yet ours is the first work, to our knowledge, to implicate CD24 in senescence. In cancer, CD24 expression is strongly associated with reduced life expectancy and is used as a diagnostic marker for patient prognosis^89^. Therefore, based on the evidence presented here, it is plausible that this role of CD24 is conserved in senescent cells, and CD24 expression may have potential as a diagnostic marker for senescence. It is important to note, however, that as the senolytic effect was exclusive to non-hematopoietic cells (Fig. 5E-G) and CD24 is expressed on many immune cell types^99^, its application to detect senescent cells will likely only apply to mesenchymal cells. The advantage compared to current senescence biomarkers (p16, p21) is that CD24 is a cell surface protein, which permits simultaneous mesenchymal cell purification alongside tracking, targeting, and sorting live senescent cells through cytometric methods. Thus, the implementation of this marker may have important implications for pre-clinical senolytic screens in murine models, and perhaps even validation of senolysis in human clinical studies. We acknowledge, however, that additional studies are needed to evaluate the potential role of CD24 in marking senescent cells in other tissues beyond the bone microenvironment and across species.

An important feature of our study is that it builds upon transcriptomic studies through investigating protein expression, which has several advantages: protein is more stable and has a longer half-life than RNA^100–103^, protein expression is more conserved than mRNA among species^104^, and mRNA transcription does not necessarily predict protein translation^102,105^. Therefore, the application of CyTOF, perhaps even in combination with scRNA-seq, allows for the rigorous investigation of senescent cells at the single-cell resolution. Importantly, our validation of CyTOF antibodies, particularly p16 and p21, provides confidence that this panel can reproducibly detect senescent cells in various applications. We expect that this established workflow can be utilized to delineate tissue-specific senescent cell identities and overcome the technical challenges of senescent cell handling and antibody specificity.

We acknowledge potential limitations of our study. Specifically, CyTOF relies on a pre-specified panel of antibodies, which limits exploration of further populations. This prevented the inclusion of additional markers, e.g. for cell identity^106–108^ and senescence^90,109,110^. We also acknowledge that there may well be further heterogeneity within our working classification of bone/marrow mesenchymal cell populations. However, in addition to being consistent with the existing literature on the identity of these cell populations, this classification does provide a useful framework for the subsequent analyses focusing on senescent cells by at least providing a provisional identity to these cells. We sought to address this limitation with the inclusion of scRNA-seq analysis, and we observed similarly clustered populations with both analyses. Nonetheless, it will be of interest to define newly discovered senescence markers and cell types, particularly those difficult to isolate in suspension (e.g. deeply embedded osteocytes), as our understanding of the aging bone microenvironment advances further. Finally, it is important to acknowledge that our characterization of p16KB and p21KB senescent cells applies only to mesenchymal cells; whether “senescent” immune cells exhibit similar characteristics or not remains to be defined.

In summary, we provide a new approach to define senescent cells *in vivo* at the single cell level using multiplexed protein profiling by CyTOF. Importantly, our definition of senescent cells is entirely consistent with a recent consensus from the ICSA^45^ and includes p16 or p21 positivity, growth arrest, upregulation of anti-apoptosis pathways, expression of a SASP, evidence of DNA damage, and virtual absence in youth with a marked increase with aging – all of which we defined within the same cells. From a translational perspective, we provide a deeper characterization of specific cell types targeted by senolytic clearance, including identifying a specific CD24+ osteolineage population, thereby facilitating the ultimate goal of defining better targets and approaches for the treatment of osteoporosis. In addition, the unique property of BCL-2 to identify true aging-specific senescent cells strongly support the development of new BCL-2 (and perhaps other BCL-related protein) inhibitors with optimized side-effect profiles to specifically target aged senescent cells for clearance. Finally, our work also points to the potential utility of CD24 for the identification and tracking of senescent cells in the bone microenvironment and perhaps in other tissues as a tool to evaluate the efficacy of senolytic treatments.

## Supporting information

Doolittle et al. Supplementary Figures and Tables

## ACKNOWLEDGEMENTS

We would like to thank the Mayo Clinic Immune Monitoring Core, and specifically Samera Farwana, for the efforts in planning and running CyTOF samples.

## AUTHOR CONTRIBUTIONS

M.L.D. and S.K. conceived and directed the project. M.L.D. and S.K. designed the experiments with input from K.P., D.G.M., and J.N.F. Experiments were performed by M.L.D., J.L.R., S.J.V., and J.K. Data was analyzed by M.L.D., and interpreted with input from S.K. and D.S. M.L.D. and S.K. wrote the manuscript, which all authors then reviewed. S.K. supervised all experimental design, data analyses, and manuscript preparation.

## Key Resources Table

| **Key Resources Table** |  |  |
| --- | --- | --- |
| **Reagent or resource** | **Source or reference** | **Identifiers** |
| **CyTOF Antibodies** |  |  |
| CD45 (30-F11) | Fluidigm | 3089005B |
| CD146 (ME-9F1) | Fluidigm | 3141016B |
| LeptinR (Goat polyclonal) | R&D Systems | AF497 |
| Nestin (Rat-401) | Biolegend | 655102 |
| Ly-6A/Sca-1 (D7) | Fluidigm | 3169015B |
| CD24 (M1/69) | Biolegend | 101802 |
| CD140a / PDGFRα (APA5) | BioLegend | 135902 |
| SDF-1 / CXCL12 (79018) | R&D Systems | MAB350-100 |
| Ostelectin/Itga11 (Rabbit polyclonal) | Abcam | ab198826 |
| CD200 (OX-90) | Biolegend | 123802 |
| CD29 (HMβ1-1) | Biolegend | 102235 |
| Runx2 (2B9) | Abcam | ab76956 |
| SP7 (Rabbit polyclonal) | Invitrogen | PA5-40411 |
| ALPL (Goat polyclonal) | R&D Systems | af2910 |
| OCN (E-6) | Santa Cruz | sc-376835 |
| E11/Podoplanin (8.1.1) | BioLegend | 127401 |
| Dmp1 (Sheep polyclonal) | ThermoFisher | PA5-47621 |
| Sclerostin (Rabbit polyclonal) | Abcam | ab63097 |
| PPARγ (Rabbit polyclonal) | Invitrogen | PA3-821A |
| Adiponectin (11H4L4) | Invitrogen | PA1-84881 |
| FLAG (L5) | Biolegend | 637301 |
| p16 (EPR20418) | Abcam | ab232402 |
| p21 (F-5) | Santa Cruz | sc-6246 |
| p53 (EPR20416-124) | Abcam | ab252388 |
| CENP-B (F-4) | Santa Cruz | sc-376283 |
| MCP-1 (2D8) | Thermo Fisher | MA5-17040 |
| TNFα (MP6-XT22) | Fluidigm | 3162002B |
| PAI-1 (1D5) | Abcam | ab125687 |
| IL-6 (MP5-20F3) | Fluidigm | 3167003B |
| IL-1α (ALF-161) | Biolegend | 503202 |
| IL-1β (D6D6T) | Cell Signaling | 31202 |
| CXCL1 (48415) | R&D Systems | MAB453-500 |
| pNFkB (93H1) | Cell Signaling | 3033 |
| pATM (EPR895) | Abcam | ab217838 |
| BCL-2 (BCL/10C4) | Biologend | 633502 |
| Ki67 (B56) | Fluidigm | 3168007B |
| **Cell Sorting Reagents** |  |  |
| CD24-PE/Cyanine7 | Biologend | 101821 |
| CD45-FITC | Biologend | 103107 |
| SYTOX Blue | ThermoFisher | S34857 |
| Lineage Depletion Kit (mouse) | Miltenyi Biotec | 130-090-858 |
| **Chemicals, Peptides, and Recombinant Proteins** |  |  |
| AP20187 | MedChemExpress | HY-13992 |
| Dasatinib | LC Laboratories | D-3307 |
| Quercetin | Sigma-Aldrich | Q4951 |
| Collagenase | Sigma-Aldrich | C9891 |
| Liberase DL | Sigma-Aldrich | 5401160001 |
| 1X RBC Lysis Buffer | ThermoFisher | 00-4333-57 |
| DPBS | ThermoFisher | 14190144 |
| DMEM | ThermoFisher | 11885-076 |
| MEM $\alpha$ | ThermoFisher | A1049001 |
| Opti-MEM | ThermoFisher | 31985062 |
| Fetal Bovine Serum (FBS) | Gemini Bio-Products | GEM100-106 |
| Antibiotic-Antimycotic | ThermoFisher | 15240096 |
| Gentamicin | Sigma-Aldrich | G1397 |
| Ascorbic Acid | Sigma-Aldrich | A8960 |
| Beta-Glycerophosphate | Sigma-Aldrich | G9422 |
| Bovine Serum Albumin | Sigma-Aldrich | A7906 |
| Alizarin Red | Millipore | TMS008C |
| 1-Step NBT/BCIP Substrate Solution (ALP) | ThermoFisher | 34042 |
| Crystal Violet | Sigma-Aldrich | C0775 |
| DAPI ProLong Diamond Antifade Mountant | ThermoFisher | P36961 |
| FuGENE 6 Transfection Reagent | Promega | E2691 |
| Trypsin-EDTA (0.5%), no phenol red | Gibco | 15400054 |
| Etoposide | Sigma-Aldrich | 341205 |
| DMSO | Sigma-Aldrich | D8418 |
| QIAzol Lysis reagent | Qiagen | 79306 |
| **Critical Commercial Assays** |  |  |
| Maxpar X8 Antibody Labeling Kit | Fluidigm | 201149A |
| Cell-ID 20-Plex Pd Barcoding Kit | Fluidigm | 201060 |
| Cell-ID Intercalator-Ir | Fluidigm | 201192B |
| Cell-ID Intercalator-Rh | Fluidigm | 201103A |
| EQ Four Element Calibration Beads | Fluidigm | 201078 |
| Senescence β-Galactosidase Staining Kit | Cell Signaling Technology | 9860 |
| Chromium Next GEM Single Cell 3' GEM, Library & Gel Bead Kit v3.1 | 10X Genomics | PN-1000128 |
| Qubit dsDNA HS and BR Assay Kits | ThermoFisher | Q32851 |
| **Software and algorithms** |  |  |
| Cytobank | Cytobank Inc. | https://premium.cytobank.org |
| R v4.03 | R Project for Statistical Computing | https://www.r-project.org |
| Seurat v4.0 | <sup>111</sup> | https://satijalab.org/seurat |
| Nebulosa v3.13 | <sup>112</sup> | https://github.com/powellgenomicslab/Nebulosa |
| CytoTree v1.03 | <sup>113</sup> | https://github.com/JhuangLab/CytoTree |
| GraphPad Prism 8 | GraphPad | http://www.graphpad.com/scientific-software/prism |
| ImageJ | NIH | https://imagej.nih.gov/ij |
| **Recombinant DNA** |  |  |
| pcDNA-5/TO-p16ink4a | This paper | N/A |
| pCMV6-Cdkn1a | Origene | MR227529 |
| pCMV6-Bcl-2 | Origene | MR226785 |
| pCMV6-Bglap (Ocn) | Origene | MR226351 |
| pCMV6-Dmp1 | Origene | MR221826 |
| pCMV6-Sost | Origene | MR222588 |
| pCMV6-Pdpm | Origene | MR201468 |
| **Experimental Models: Organisms/Strains/Cell lines** |  |  |
| INK-ATTAC | <sup>12</sup> | N/A |
| C57BL/6N | Charles River Laboratories | RRID:IMSR_CRL:027 |
| Alb.B6.Cdkn2a-Luciferase ($p16^{Luc}$) | <sup>36</sup> | RRID:IMSR_NCIMR:01XBT |
| U2OS Cell Line | ATCC | ATCC (HTB-96); RRID:CVCL_0042 |

## Resource Availability

### Lead contact

Further information and requests for resources and reagents should be directed to and will be fulfilled by Dr. Sundeep Khosla.

### Materials availability

All unique/stable reagents generated in this study will be freely available from the lead contact to academic researchers with a completed Materials Transfer Agreement.

#### Animals

All animal studies were performed under protocols approved by the Institutional Animal Care and Use Committee (IACUC), and experiments were performed in accordance with Mayo Clinic IACUC guidelines. Mice were housed in ventilated cages and maintained within a pathogen-free, accredited facility under a twelve-hour light/dark cycle with constant temperature (23°C) and access to food and water ad libitum. Mice used included *INK-ATTAC* ^12^, *C57BL/6N* Wild Type (WT), and p16 luciferase reporter mice (*p16^Luc^*) ^36^. Two ages were used for all studies: Young-6 month, and old-24 month. All young mice were untreated. Old *INK-ATTAC* mice were randomized by weight for treatment with either vehicle (4%ETOH 10%PEG-400 and 2%Tween) or 10mg/kg AP dissolved in vehicle, administered subcutaneously twice weekly for two weeks. Old C57BL/6N mice were randomized by weight for treatment with vehicle (10% EtOH, 30% PEG-400, 60% Phosal-50) or Dasatinib + Quercetin (D+Q) (Dasatinib 5mg/kg and Quercetin 50mg/kg) dissolved in vehicle, administered by oral gavage for three consecutive days and harvested at two weeks post-treatment. Each cohort consisted of the following groups: *INK-ATTAC:* Young (n=15; 10 female, 5 male), Old + Vehicle (n=12; 6 female, 6 male), Old + AP (n=13; 6 female, 7 male). C57BL/6N: Young (n=17; 10 female, 7 male), Old + Vehicle (n=15; 8 female, 7 male), Old + D+Q (n=16; 10 female, 8 male). In comparisons of young versus old mice, Old + Vehicle mice were used, for reduction purposes. To our knowledge, there are no effects of vehicle administration for either treatment on aging, senescent, or skeletal outcomes. For CD24 cell isolation and scRNA-seq studies, 24-27-month untreated male and female INK-ATTAC mice were used, with sexes indicated in figure legends per experiments.

#### Consideration of sex as a biological variable

Per NIH guidelines ^114,115^, we studied both female and male mice. In order to test for possible effects of sex on our primary endpoints ^116^, we performed 2-way ANOVA tests on several important parameters of aging and senolytic treatment (Supplementary Table 1). We found that neither sex alone nor interaction between sex and age was significant, indicating that these cellular effects of aging or senolytic treatment are not dependent on sex. Thus, both males and females were analyzed together.

#### Dissociation and purification of mesenchymal cells from skeletal tissue

Mice were euthanized according to standardized and approved IACUC protocols. Femurs and tibiae were isolated, cleaned of soft tissue, cut at both ends, and marrow centrifuged out of the diaphyses and metaphyses into a collection tube. Marrow was resuspended in 1mg/mL Liberase DL (Sigma) diluted in FACS buffer (0.5% BSA [Sigma] in PBS) and digested at 37°C for 30 minutes to increase yield of stromal cells released from the vasculature fraction as previously described ^33^. Diaphyses and metaphyses cleared of bone marrow were gently crushed, rinsed in PBS, and then digested in 300 Units/mL of Collagenase IA (Sigma), diluted in MEM α (ThermoFisher), 3 times for 25 minutes each. Bone and marrow solutions were then combined and treated with RBC lysis buffer (ThermoFisher) to clear erythrocytes. The sample was then depleted of cells expressing hematopoietic lineage markers (CD5, CD45R [B220], CD11b, Gr-1 [Ly-6G/C], 7-4, and Ter-119) using Magnet Assisted Cell Sorting (MACS) and the Lineage Cell Depletion Kit (Miltenyl Biotec).

#### Cell culture

Primary mouse BMSCs were generated from 6-month-old C57BL/6N mice by digesting freshly dissected femurs and tibias 3 times for 25 minutes each, followed by RBC lysis, as described above. After expansion in growth media (DMEM [ThermoFisher] + 15% FBS [Gemini Bio-Products] + 1X Antibiotic/Antimycotic [ThermoFisher] + 1X Gentamicin [Sigma]) in hypoxic condition (2% O2), BMSCs were seeded at 4×10^4^ cells/cm^2^ in 75cm^2^ flasks and treated for 48 hours with either vehicle (0.1% DMSO [Sigma]) or 20uM of etoposide (MilliporeSigma, St. Lous, MO) dissolved in vehicle, followed by maintenance in growth media for 6 days. Cells were then dissociated using Trypsin-EDTA (Gibco) and processed for CyTOF. CD24- and + cells were maintained in the same growth media, as described above. For osteogenesis experiments, CD24- and + cells were seeded at 2×10^4^ cells/cm^2^ in 96-well plates, upon which media was changed to osteogenic medium (MEM α + 10% FBS + 1X Anti/Anti + 10mM β-Glycerophosphate [Sigma] + 50mg/ml Ascorbic Acid [Sigma]) and differentiated until the indicated time point, changing media every 48 hours. U2OS cells were maintained in normoxic conditions (5% O_2_) in 75cm^2^ flasks in DMEM + 10% FBS + 1X Antibiotic/Antimycotic and seeded at 2×10^4^ cells/cm^2^ in 6-well plates for transfection. Expression constructs were transfected into U2OS cells using 1.5μg of DNA and 4.5μL FuGENE 6 Transfection Reagent (Promega) per well. DNA-FuGENE mixtures were combined in Opti-MEM (ThermoFisher), then added dropwise to U2OS cells. After 24 hours, cells were lifted using Trypsin-EDTA and processed for CyTOF, while separately lysing 10% of each cell sample in QIAzol for qPCR validation.

#### p16 expression vector construction

The pcDNA5/TO-p16ink4a plasmid was constructed by cloning the open reading frame of mouse *p16^Ink4a^* into the BamHI site of pcDNA5/TO (Promega, Madison, WI).

#### CyTOF processing

Custom conjugated antibodies were generated in-house through the Mayo Clinic Hybridoma Core using Maxpar X8 Ab labeling kits (Fluidigm) according to the manufacturer’s protocol. Isolated cells were resuspended in 1 mL of Cell Staining Buffer (CSB) (Fluidigm) and incubated for 5 minutes with 0.5 μm Cisplatin solution (Fluidigm) in PBS. Samples were then washed twice with CSB. An antibody cocktail of the entire phenotyping panel was prepared as a master mix prior to adding 50 μL of cocktail to samples resuspended in 50 μL of CSB. Samples were then incubated at room temperature for 45 minutes. Samples were washed twice then fixed with 2% PFA (Fluidigm) in PBS. After fixation and wash, samples were resuspended in 30 nM intercalation solution (Fluidigm) and incubated overnight at 4°C. On the following morning, cells were washed with PBS and resuspended in a 1:10 solution of calibration beads and cell acquisition solution (CAS) (Fluidigm) at a concentration of 0.5×10^6^ cells/mL. Prior to data acquisition, samples were filtered through a 35 μm blue cap tube (Falcon). The sample was loaded onto a Helios CyTOF system (Fluidigm) and acquired at a rate of 200-400 events per second. Data were collected as .FCS files using the Cytof software (Version 6.7.1014). After acquisition, intra-file signal drift was normalized to acquired calibration bead signal using Cytof software.

#### CyTOF data analysis

##### Initial processing and clustering

Cleanup of cell debris–including removal of beads, dead cells, and doublets–and negative selection of CD45+ cells was performed (Extended Data Fig. 2A) using Cytobank software ^117,118^. Visual representation of CD45-single-cell data was achieved using UMAP ^119^ (15 neighbors, 0.01 minimum distance, outliers collapsed), and viSNE mapping (5,000 iterations, 100 perplexity, 0.5 theta), the latter of which is based on the t-Distributed Stochastic Neighbor Embedding (t-SNE) algorithm ^120^. FlowSOM clustering was performed within Cytobank (hierarchical consensus, 10 iterations) and cluster labels were assigned using established literature on skeletal cell types (Table 1), with relative marker intensities per cluster visualized by heatmap. FCS files were exported, concatenated in R, then re-uploaded for visualization of merged populations. Quantified values were exported to Graphpad Prism 8 to construct plots and perform statistical analyses.

##### CITRUS analysis

CITRUS analyses ^121^ were performed in Cytobank using Significance Analysis of Microarrays (SAM) correlative association model. Nearest Shrunken Centroid (PAMR) and L1-Penalized Regression (LASSO via GLMNET) predictive association models were run simultaneously to analyze model error rates to confirm validity of the statistical model. For CITRUS assessment of median expression changes, cells were clustered by identification markers and statistics channels included all functional markers; for assessment of abundances, all markers were used for clustering. All CITRUS analyses used the following settings: 2,000 events samples per file, 2% minimum cluster size, 5 cross validation folds, and 5% false discovery rate (FDR).

##### Pseudotime

Exported FCS files from Cytobank were imported into R using the CytoTree package ^113^. Briefly, samples were merged per condition and clustered into minimum spanning trees for cell trajectory inference. Pseudotime calculation was performed by defining root clusters based on expression of stem cell markers (CD146, Sca-1, PDGFRα, Nestin, LeptinR). Representation of pseudotime was performed using diffusion mapping, along with density and trajectory plots.

#### Fluorescence-assisted cell sorting (FACS)

Single-cell suspensions of mesenchymal skeletal cells were prepared, as described above. Samples were incubated with anti-mouse CD45-FITC (and CD24-PECy7 for CD24- and + cell isolation) at 1:400 dilution in FACS buffer at 4°C for 20 minutes in the dark. Cells were then incubated with SYTOX blue at 1:4,000 for 5 minutes, spun down at 300xg for 5 minutes at 4°C, then resuspended in FACS buffer at 1×10^7^ cells/mL and analyzed on a FACS Aria II (BD Biosciences). Unstained and single-color-stained controls were used for compensation and to control the gating strategy. Post-run flow cytometry data was analyzed and visualized with Cytobank software.

#### Cell staining

##### SA-β-Gal

To assess senescence *in vitro*, cellular SA-β-Gal activity was measured as described previously ^95^ using the Cell Signaling Technology Senescence β-Galactosidase Staining Kit. Briefly, CD24+ or – cells were seeded on 8-well chamber slides at 1×10^4^ cells/cm^2^ and allowed to grow for 14 days. Cells were then washed in PBS (pH 7.4) and fixed with 1X fixative solution for 5 min, then washed three times using PBS. Cells were then incubated in 1X SA-β-Gal staining solution at 37 °C for 16 hr. Cells were washed in ice-cold PBS and mounted with DAPI ProLong (ThermoFisher) staining nuclei for cell counting. In blinded fashion, ten images per well were taken from random fields using fluorescence microscopy (Nikon Eclipse Ti) and SA-β-Gal-positive cells were counted and reported as a percentage of total cells.

##### Crystal violet

CD24- and + cells were seeded directly from FACS-mediated isolation into 25cm^2^ flasks and allowed to grow for 7 days. Cells were then washed in PBS and fixed in 4% PFA for 20 min. Fixed cells were washed again with PBS and stained with 1% crystal violet in 20 % ethanol for 20 min. Excess dye was removed by washing with distilled water (dH_2_0) and images were acquired upon drying. Colony forming efficiency (CFE) was determined for each sample by counting colonies containing over 50 cells, then dividing by total sorted cells.

##### Alizarin Red and Alkaline Phosphatase

Cells were washed with PBS then fixed with 4% PFA for 10 minutes. For Alkaline Phosphatase (ALP) analysis, the fixed cells were stained with 1-Step NBT/BCIP Substrate Solution (ThermoFisher) in the dark for 30 minutes. To detect mineralization, cells were fixed in 4% PFA and stained with Alizarin Red (Millipore) for 30 minutes; both stains were washed with dH_2_0 and let dry before imaging.

#### Quantitative real-time polymerase chain reaction (qPCR) analysis

Total RNA was extracted according to the manufacturer’s instructions using QIAzol Lysis Reagent. Purification with RNeasy Mini Columns (QIAGEN, Valencia, CA) was subsequently performed. On-column RNase-free DNase solution (QIAGEN, Valencia, CA), was applied to degrade contaminating genomic DNA. RNA quantity was assessed with Nanodrop spectrophotometry (Thermo Fisher Scientific, Wilmington, DE). Standard reverse transcriptase was performed using High-Capacity cDNA Reverse Transcription Kit (Applied Biosystems by Life Technologies, Foster City, CA). Transcript mRNA levels were determined by qRT-PCR on the ABI Prism 7900HT Real Time System (Applied Biosystems, Carlsbad, CA), using SYBR green (Qiagen, Valencia, CA). The mouse primer sequences, designed using Primer Express Software Version 3.0 (Applied Biosystems), for the genes measured by SYBR green are provided in Supplementary Table 2. Input RNA was normalized using two reference genes (*Actb, Gapdh*) from which the most stable reference gene was determined by the geNorm algorithm. For each sample, the median cycle threshold (Ct) of each gene (run in triplicate) was normalized to the geometric mean of the median Ct of the most stable reference gene. The delta Ct for each gene was used to calculate the relative mRNA expression changes for each sample. Genes with Ct values > 35 were considered not expressed (NE), as done previously ^6^.

#### scRNA-seq library preparation

Live Lin-CD45-cells, isolated by FACS, were washed twice in 1x PBS + 0.04% BSA and immediately submitted to the Mayo Clinic Genome Analysis Core for single-cell sorting. The cells were counted and measured for viability using the Vi-Cell XR Cell Viability Analyzer (Beckman-Coulter). The Chromium Next GEM Single Cell 3’ Library and Gel Bead Kit (10x Genomics) was used for cDNA synthesis and standard Illumina sequencing primers and a set of unique i7 Sample dual indices (10x Genomics) were added to each cDNA pool. All cDNA pools and resulting libraries were measured using Qubit High Sensitivity assays (Thermo Fisher Scientific), Agilent Bioanalyzer High Sensitivity chips (Agilent) and Kapa DNA Quantification reagents (Kapa Biosystems). Libraries were sequenced at 50,000 fragment reads per cell following Illumina’s standard protocol using the Illumina cBot and HiSeq 3000/4000 PE Cluster Kit. The flow cells were sequenced as 100 X 2 paired end reads on an Illumina HiSeq 4000 using HiSeq 3000/4000 sequencing kit and HCS v3.3.52 collection software. Base-calling was performed using Illumina’s RTA version 2.7.3. 10X Genomics Cell Ranger Single Cell Software Suite (v6.0.0) was used to demultiplex raw base call (BCL) files generated from the sequencer into FASTQ files. The pipeline input FASTQ files for each sample to perform alignment to the reference genome, filtering, barcode counting and UMI counting.

#### scRNA-seq analysis

Seurat package (v4.0) ^111^ was used in R to perform integrated analyses of single cells. Genes expressed in < 3 cells and cells that expressed < 200 genes and >20% mitochondria genes were excluded for downstream analysis in each sample. Each dataset was SCTransform-normalized and the top 3000 Highly Variable Genes (HVGs) across cells were selected. The datasets were integrated based on anchors identified between datasets before Principal Component Analysis (PCA) was performed for linear dimensional reduction. Shared Nearest Neighbor (SNN) Graph were constructed to identify clusters on the low-dimensional space (top 30 statistically significant principal components (PCs). Enriched marker genes in each cluster conserved across all samples were identified. An unbiased clustering according to the recommendations of the Seurat package was used, and a resolution of 0.8 led to 12 distinct cellular clusters (Extended Data Fig. 8). For Uniform Manifold Approximation and Projection for Dimension Reduction (UMAP) calculations, the RunUMAP function (dims = 1:40, reduction = “pca”) was utilized, and both DimPlot (Seurat) and plot_density (Nebulosa) used for plotting.

#### Statistics

In-text results and bar plots are mean ± standard deviation. Graphical data represented as box plots show median and interquartile range, and error bars represent minimum and maximum values. Sample sizes were determined based on previously conducted and published experiments ^16,95^ in which statistically significant differences were observed among various senescence and skeletal parameters in response to aging or senolytic treatment. Animal numbers are indicated in the figure legends, and all samples presented represent biological replicates. We did not exclude mice, samples, or data points from analyses. Non-Gaussian distributions were detected using the Shapiro-Wilk normality test. If the normality or equal variance assumptions for parametric analysis methods were not met (Shapiro-Wilk p<0.05), data were analyzed using non-parametric tests (e.g., Mann-Whitney U test). For parametric tests, differences between groups were analyzed by t-test (followed by Hold-Sidak correction with multiple comparisons) or ANOVA, where justified as appropriate; figure legends indicate the statistical tests used in each experiment. Statistical analyses were performed using GraphPad Prism (Version 8.0). A p-value <0.05 was considered statistically significant. Heatmap values were transformed by subtraction of row mean and dividing by standard deviation, visualized in Morpheus – Broad Institute. Experimental design diagrams and schematics were made using BioRender.com. Venn diagrams were made using Meta-Chart.com.

