## Supplementary material for "Multiparametric senescent cell phenotyping reveals CD24 osteolineage cells as targets of senolytic therapy in the aged murine skeleton": Doolittle et al. Supplementary Figures and Tables

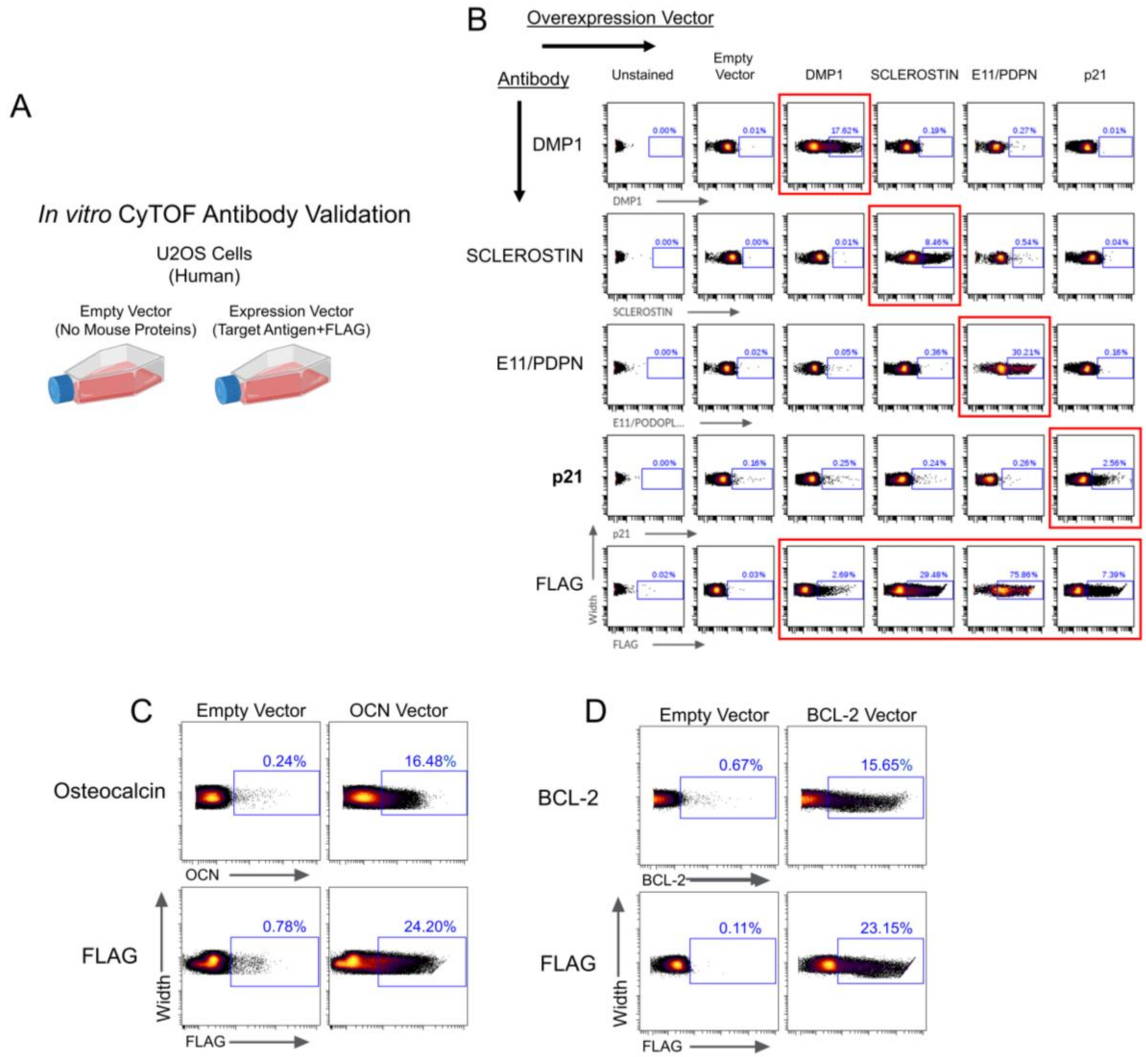


**Extended Data Figure 1. Validation of antibodies in CyTOF panel.** (A) Experimental workflow of single mouse protein expression in U2OS cells for the testing of CyTOF antibodies; (B) CyTOF plots of several expression tests run simultaneously. Each column is an individual sample, and each row is the antibody being tested. Red boxes demonstrate positive results, where signal is observed in the positive gate (blue box) and not observed in any other channel or sample. FLAG demonstrates successful expression of the DNA plasmid; (C) Additional validations of osteocalcin and BCL-2 antibodies by FLAG-tagged expression vectors.

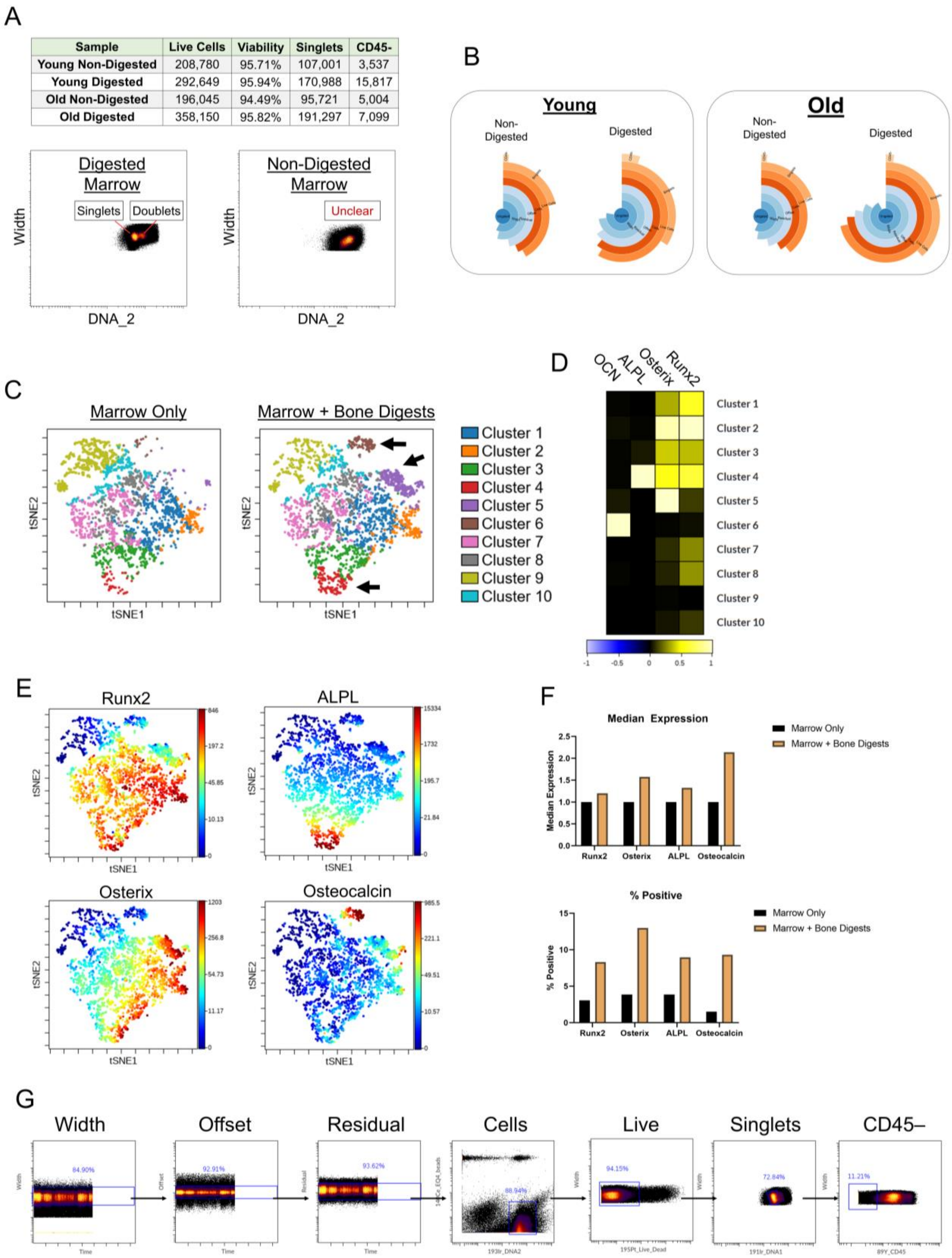


**Extended Data Figure 2. Optimization of bone and marrow digestion protocols.** (A) Quantification of results and CyTOF plots demonstrating better resolution of singlets and doublets with Liberase digestion (X-axis is DNA content, while Y-axis is width). (B) Sunburst plots from experiments trialing the Liberase digestion of bone marrow samples, demonstrating an increase in yield of total and CD45- cells, with a preservation in cell viability; (C) t-SNE plots of samples containing cells from either marrow-only or marrow combined with 3 bone digestions. FlowSOM-clustered cell populations demonstrate an emergence of several clusters, marked by arrows, when adding in cells isolated from digested bone; (D) Heatmap of median expression and (E) feature plots of osteolineage markers across clusters, demonstrating expression of ALPL, Osterix, and Osteocalcin (OCN) in emerging clusters 4, 5, and 6, respectively; (F) Overall median expression and percent-positive values of osteolineage markers, demonstrating an enrichment with cells obtained from digested bone. n=1 mouse per condition. (G) Gating strategy for cleanup and purification of mesenchymal cells from digested bone/bone marrow cell suspensions.

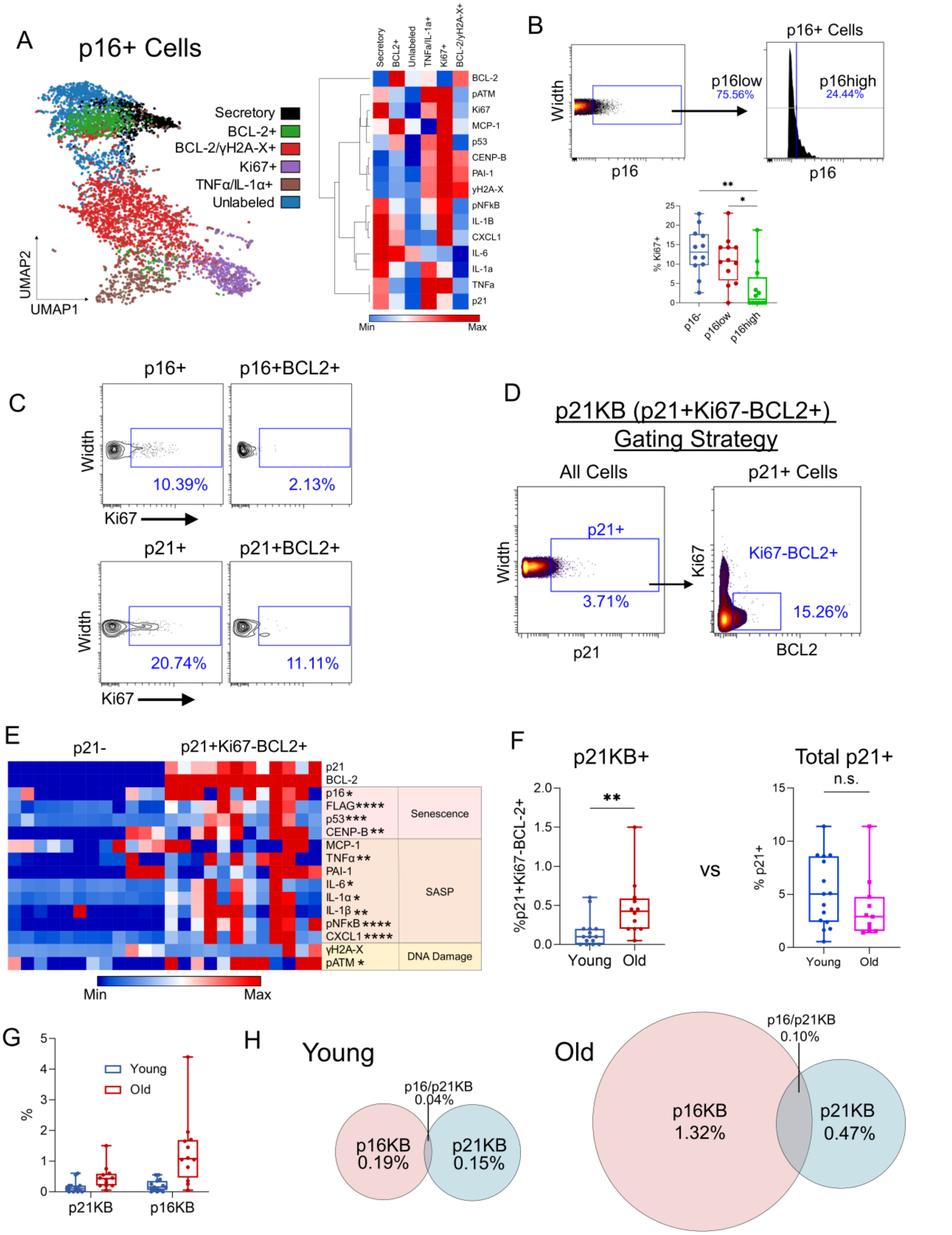


23 **Extended Data Figure 3: Defining growth-arrested subsets of p16+ and p21+ cells**

**associated with age.** (A) UMAP visualization and FlowSOM clustering of p16+ cells from young and old mice (merged) with corresponding heatmap demonstrating senescence panel marker expression in the 6 identified clusters; (B) Gating strategy for p16low and p16high cells, with %Ki67+ for each population below alongside p16- cells. (C) Dot plots of BCL2+ subsets of p16+ or p21+ cells demonstrating %Ki67+ cells in each population (blue gate); (D) Gating strategy for p21KB cells; (E) Heatmap representation of protein expression between p21KB cells and p21- cells, with asterisks indicating significance. (F) Quantification of p21KB cells with age and comparison to total p21+ cells. (G) Comparison of p21KB vs p16KB cell proportions. (H) Venn diagrams of p16KB versus p21KB cells demonstrating very few shared cells between populations. *p<0.05, **p<0.01, ***p<0.001, ****p<0.0001. Mann-Whitney test.

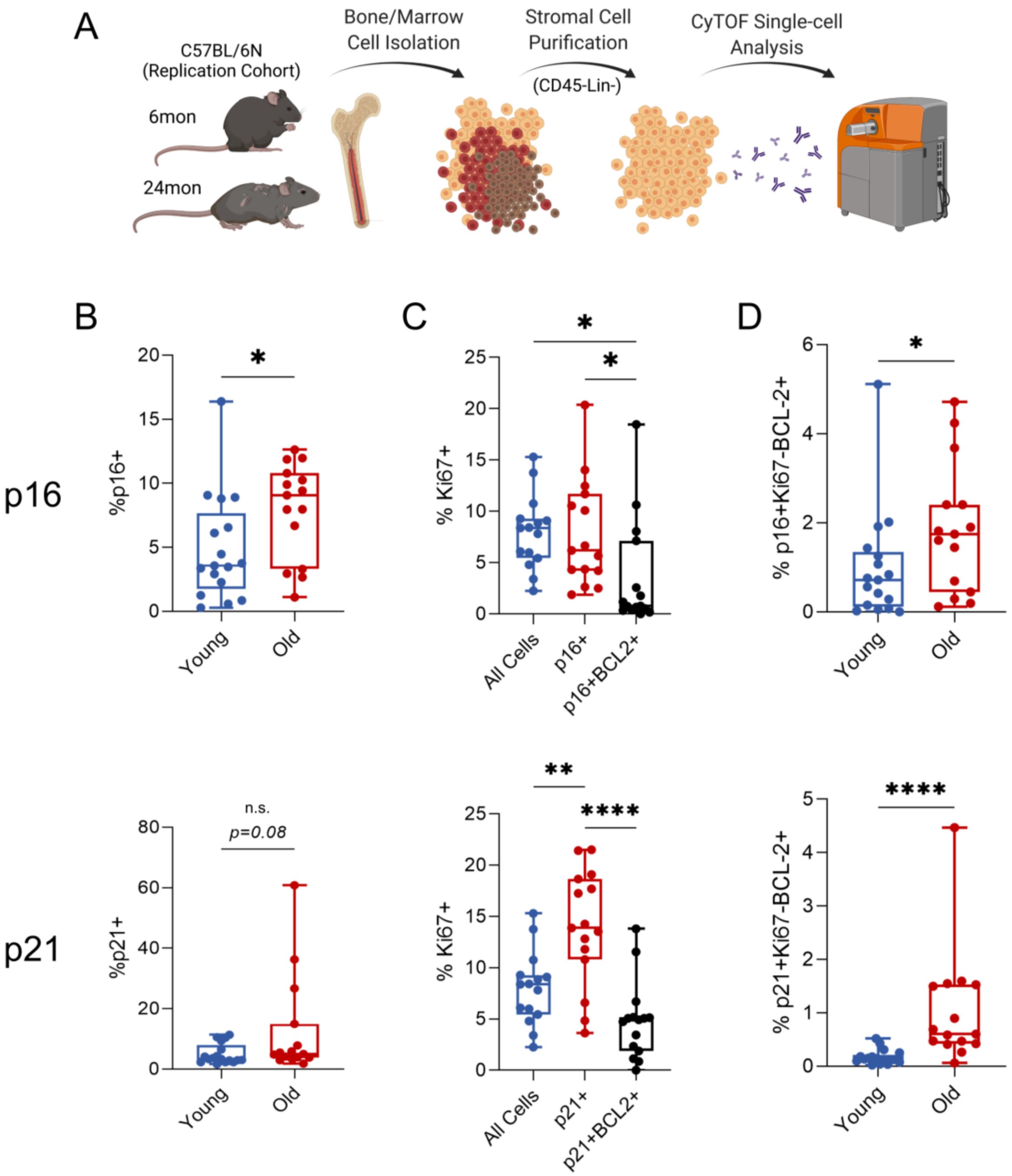


**Extended Data Figure 4. Investigation and replication of BCL-2 results.** (A) Schematic of replication cohort workflow, containing n=17 young and n=15 old C57BL/6N mice (Old + Vehicle mice from D+Q experiment); (B) Percent-positive plots of p16+ p21+ cells between young and old mice. (C) % Ki67+ in all cells, p16+/p21+ cells, and BCL-2+ subsets of each. (D) %p16KB or p21KB cells with age. *p<0.05, **p<0.01; (B, D) Unpaired t test or Mann-Whitney test as appropriate. (C) Multiple t tests with Holm-Sidak Correction.

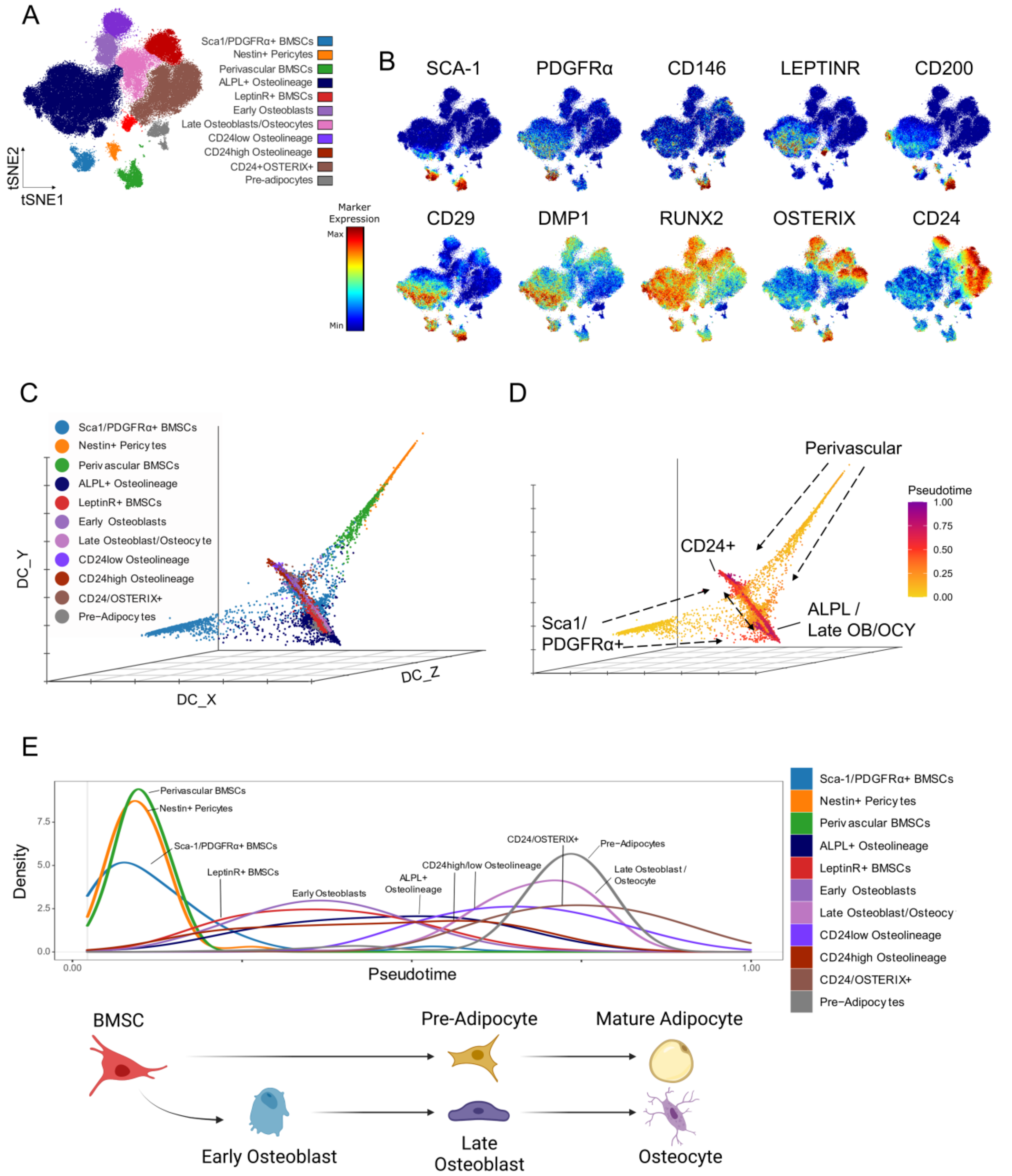


**Extended Data Figure 5. Single-cell CyTOF reconstruction of the mesenchymal bone/bone marrow microenvironment.** (B) t-SNE visualization and FlowSOM clustering of CD45-Lin- bone and marrow cells analyzed by CyTOF. (B) t-SNE feature plots depicting protein expression of cluster-defining phenotypic markers; (C) Diffusion map visualization of clustered populations; (D) Pseudotime trajectory displayed on diffusion map displaying CD24 osteolineages (CD24+), ALPL+ osteolineage cells (ALPL), and late osteoblast/osteocyte (Late OB/OCY) clusters as late in differentiation downstream of the Perivascular and Sca1/PDGFRα+ BMSC clusters. Note the divergence of the CD24+ versus the ALPL+ osteolineage and late osteoblast/osteocyte clusters. Dotted arrows depict pseudotime trajectory; (E) Density of clusters along pseudotime consistent with established cell differentiation patterns, shown by the diagram below.

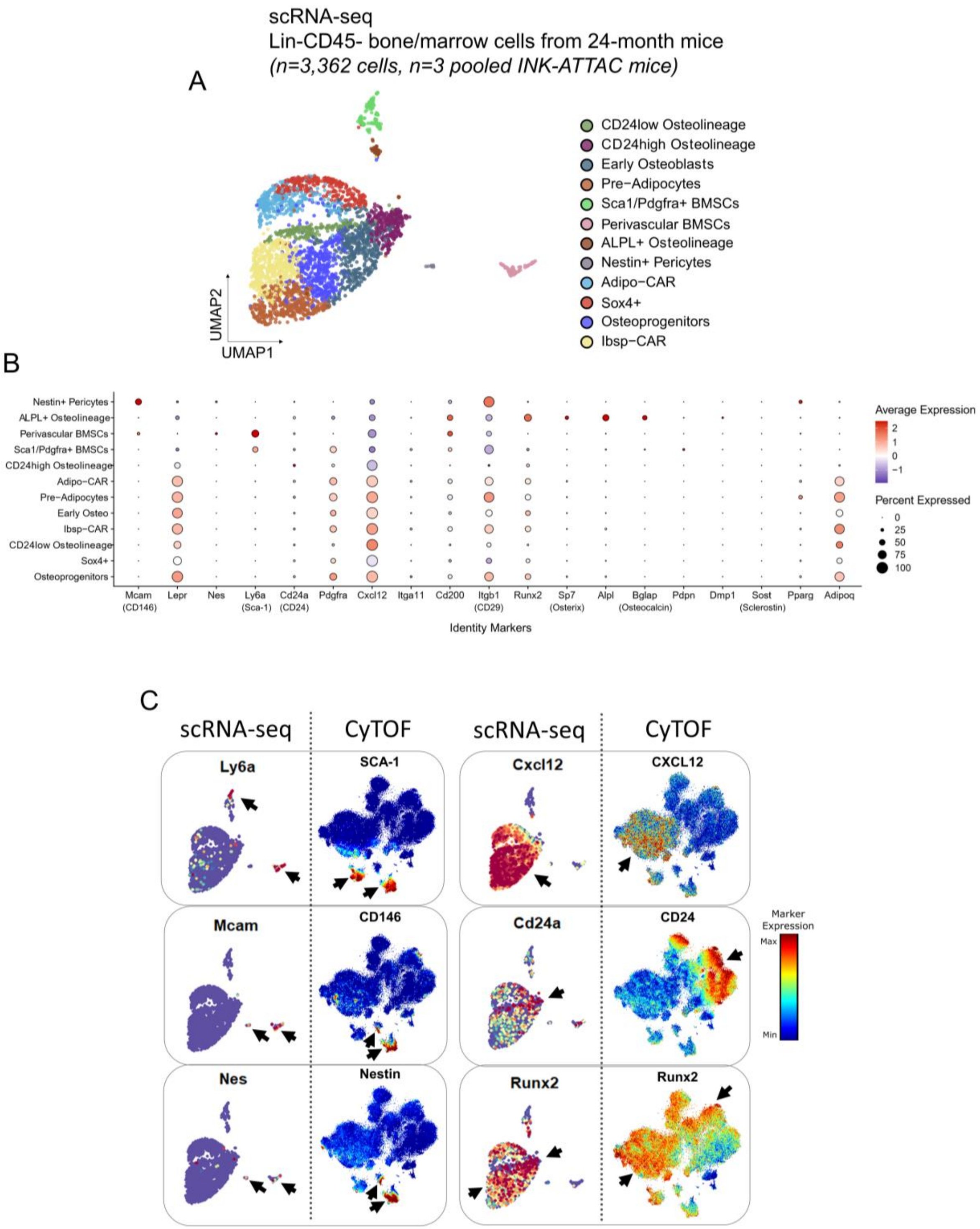


**Extended Data Figure 6. Validation of CyTOF populations by scRNA-seq.** (A) UMAP visualization of n=3,362 clustered Lin-CD45- cells from the digested bone and marrow of n=3 24-month untreated *INK-ATTAC* mice; (B) Dot plot expression of markers defining each scRNA-seq cluster; (C) Visual comparison of multidimensional scRNA-seq and CyTOF single-cell data, demonstrating similar expression patterns of common mesenchymal markers using each tool, marked by black arrows.

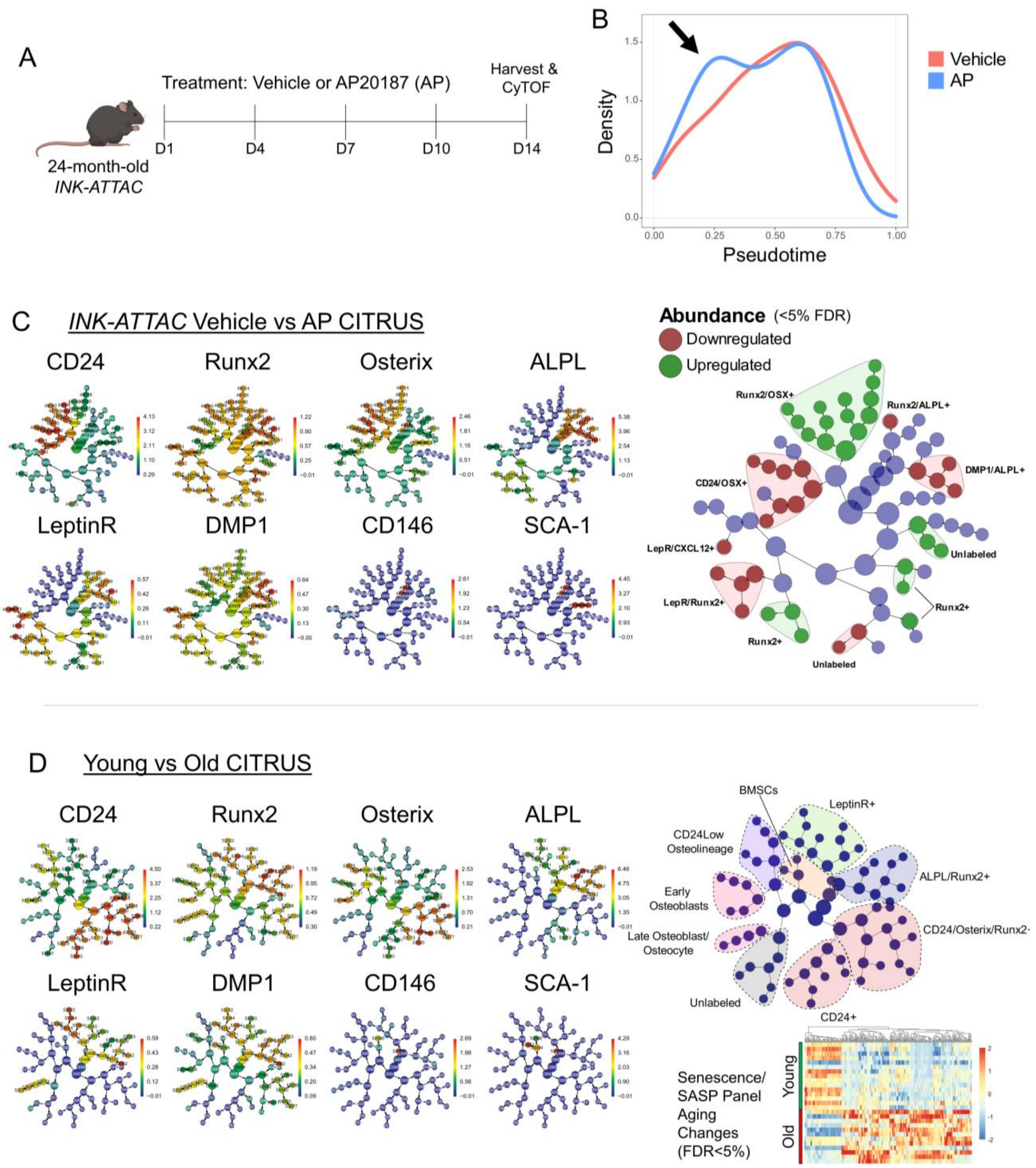


**Extended Data Figure 7. INK-ATTAC Analyses and CITRUS Defining markers.** (A) Schematic of p16+ senescent cell clearance in *INK-ATTAC* mice analyzed by CyTOF. Mice were treated with vehicle or AP every three days for 14 days (indicated by dashes); (B) Pseudotime density plot of vehicle- and AP-treated samples, demonstrating a surge in early-pseudotime cell types, marked by black arrow. (C) Expression plots for defining markers of CITRUS cluster families for vehicle- versus AP-treated *INK-ATTAC* mice. Plots demonstrating cell abundance changes within clusters after AP treatment (q<0.05), with red marking cleared clusters and green marking upregulated clusters; (D) Expression plots for defining markers of CITRUS cluster families for INK-ATTAC young (n=15) vs old (n=12) analyses corresponding to Figure 4C. Cluster families are colored and indicated by dotted lines. Heatmap demonstrates all significant (FDR<5%) CITRUS results of senescence panel changes with age: Rows are mice and each column is a cluster-marker combination that reached significance.

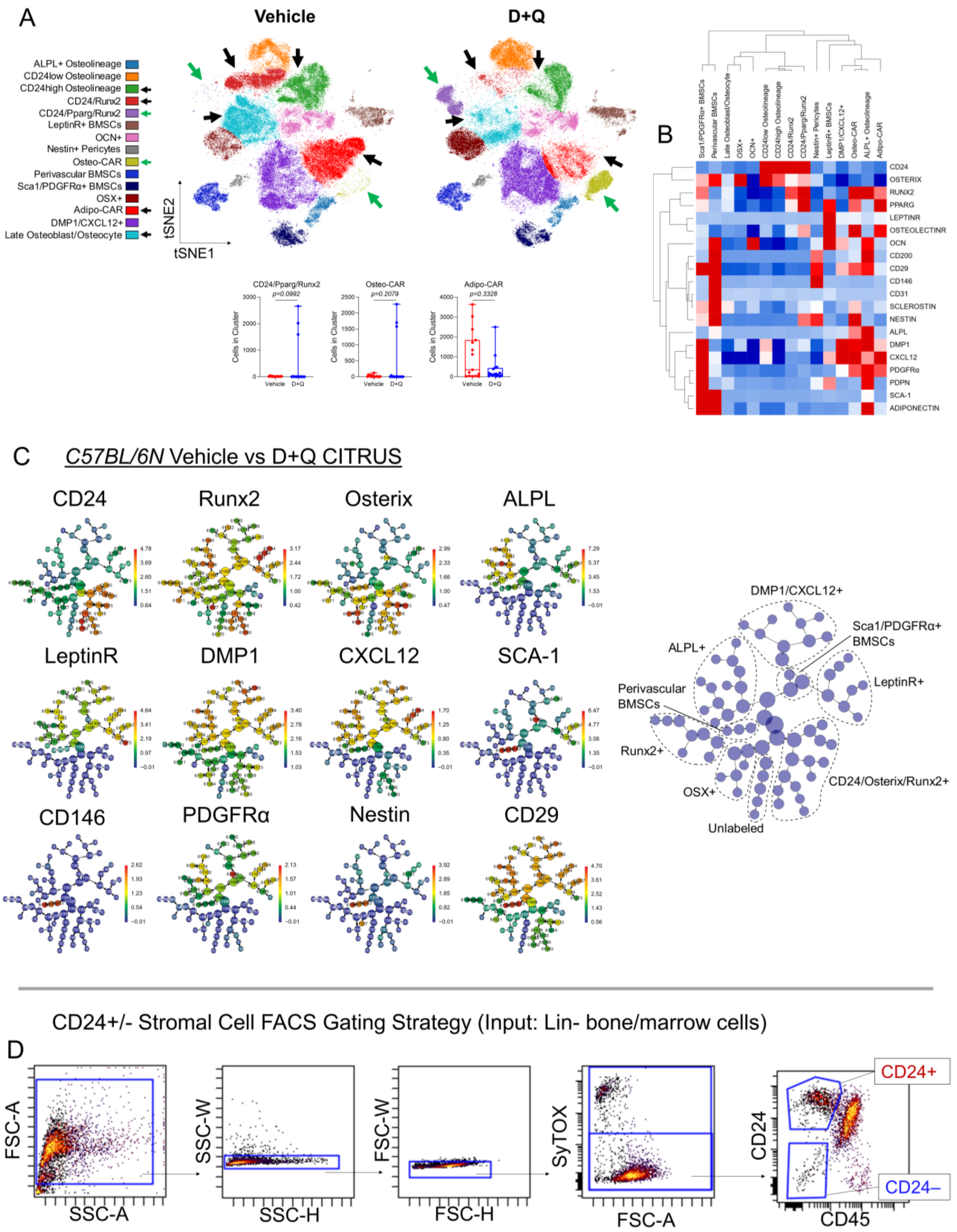


**Extended Data Figure 8. CyTOF analyses of mice undergoing D+Q Treatment.** (A) t-SNE visualization and FlowSOM clustering of bone/bone marrow cells from vehicle- or D+Q-treated old mice (n=15 vehicle, n=16 D+Q; n= 5,980 cells per mouse). Clusters visually cleared are marked by black arrows, and emerging clusters are marked by green arrows. Box plots below depict clusters marked by arrows yet were not significantly cleared; (B) Heatmap representation of the cell clusters and protein expression of identification markers; (C) Expression plots for defining markers of CITRUS cluster families for *C57BL/6N* vehicle vs D+Q analyses corresponding to Figure 5D. (C) Gating strategy for isolation of live (SYTOX-) CD45-CD24+ and – cells by FACS.

**Supplementary Table 1. Assessment of sex as a biological variable.** 2-way ANOVA results found no substantial interaction effects of sex on primary endpoints of aging and senolytic treatment.

| Metric | Source of Variation | % of total variation | P value | P value summary | Significant ? |
|---|---|---|---|---|---|
| p16 median expression | Interaction | 4.718 | 0.0846 | ns | No |
| | Sex | 0.01463 | 0.9209 | ns | No |
| | **Age** | **52.86** | **<0.0001** | ******** | **Yes** |
| % p16+ cells | Interaction | 5.05 | 0.1671 | ns | No |
| | Sex | 5.481 | 0.1508 | ns | No |
| | **Age** | **31.37** | **0.0017** | ****** | **Yes** |
| % p16KB cells | Interaction | 5.304 | 0.1346 | ns | No |
| | Sex | 8.761 | 0.0582 | ns | No |
| | **Age** | **35.29** | **0.0006** | ******* | **Yes** |
| % p16KB cells | Interaction | 2.326 | 0.4334 | ns | No |
| | Sex | 0.3027 | 0.7761 | ns | No |
| | **Senolytic (AP)** | **21.14** | **0.0253** | ***** | **Yes** |
| % CD24high osteolineage cells | Interaction | 1.439 | 0.4861 | ns | No |
| | Sex | 1.056 | 0.5502 | ns | No |
| | **Senolytic (AP)** | **37.29** | **0.0016** | ****** | **Yes** |

**Supplementary Table 2. qPCR primer sequences.**

| Gene | Forward Primer Sequence | Reverse Primer Sequence |
|---|---|---|
| *Cdkn2a (p16ink4a)* | GAACTCTTTCGGTCGTACCC | AGTTCGAATCTGCACCGTAGT |
| *Cdkn2a (p19arf)* | GGCTTTCGTGAACATGTTGTTG | AACGTTGCCCATCATCATCA |
| *Cdkn1a (p21)* | GAACATCTCAGGGCCGAAAA | TGCGCTTGGAGTGATAGAAATC |
| *Cd24a (CD24)* | TGCTTCTGGCACTGCTCCTA | CGGTGCAACAGATGTTTGGTT |
